# Sex differences in dispersal predict sex differences in helping across cooperative birds and mammals

**DOI:** 10.1101/2023.09.25.557200

**Authors:** Patrick Fenner, Thomas E. Currie, Andrew J. Young

## Abstract

Sex differences in cooperation are widespread, but their evolution remains poorly understood. Here we use phylogenetic comparative methods to test the Dispersal hypothesis for the evolution of sex differences in contributions to cooperative care across the cooperatively breeding birds and mammals. The Dispersal hypothesis predicts that, where non-breeding individuals of both sexes help to rear offspring within their natal group, the more dispersive sex will contribute less (either because leaving the natal group earlier reduces the downstream direct benefit from cooperation or because dispersal activities trade-off against cooperation). Our analyses reveal (i) support for the Dispersal hypothesis (sex biases in dispersal predict sex biases in natal cooperation across taxa), and (ii) that this pattern cannot be readily attributed to alternative hypothesized drivers of sex differences in cooperation (kin selection, heterogamety, paternity uncertainty, patterns of parental care or differences between birds and mammals). Our findings help to clarify the evolutionary drivers of sex differences in cooperation and highlight the need for single-species studies to now tease apart whether sex differences in dispersal predict sex differences in natal cooperation because dispersal impacts the direct benefits of natal cooperation (as is often proposed) or because activities that promote dispersal trade-off against natal cooperation.

## INTRODUCTION

In many animal societies males and females differ in their contributions to cooperative activities, and the evolutionary origins of such sex differences in cooperation remain poorly understood [1–7]. In many cooperatively breeding species, for example, offspring of both sexes delay dispersal from their natal groups and cooperatively contribute to the feeding of future generations of their parents’ young [2, 8, 9]. Numerous findings suggest that selection for such cooperative ‘helping behavior’ typically arises at least in part via kin selection, with helpers accruing indirect fitness benefits from enhancing the fitness of relatives [2, 8–13]. However, kin selection alone cannot readily account for the evolution of sex differences in cooperation within the natal group, as male and female helpers should not differ in their mean relatedness to recipients in this context ([14]; at least with regard to autosomal genes [15]). The evolution of such sex differences in natal cooperation therefore seems likely to be attributable instead to sex differences in the net direct fitness payoff from natal cooperation, via a sex difference in the direct benefits and/or costs of cooperation [3–6, 16–19].

One general mechanism that has been hypothesized to drive the evolution of sex differences in natal cooperation across taxa is sex differences in dispersal [3–6, 16–18]. The ‘Dispersal hypothesis’ proposes that the more dispersive sex stands to gain a lower net direct fitness payoff from helping within the natal group and should therefore help at a lower rate while within the natal group. This lower net direct fitness payoff from natal helping for the more dispersive sex could arise via two general mechanisms, acting in isolation or concert. First, as helpers of the more dispersive sex are expected to stay for less time on average within their natal group, they may stand to gain a lower downstream direct fitness *benefit* from natal helping if the accrual of this direct benefit is contingent in part upon remaining in the natal group [3, 6, 16]. For example, wherever helping increases natal group size (e.g. by improving offspring survival) and members of larger groups enjoy higher survival and/or downstream breeding success [20, 21], helpers of the more dispersive sex may gain a lower downstream direct fitness benefit from helping to augment natal group size as they are likely to leave the natal group sooner [3, 6, 16–18]. A second general mechanism that may leave the more dispersive sex standing to gain a lower net direct fitness payoff from natal cooperation is if investments in cooperation trade off against investments in activities that promote dispersal success [4, 5]. For example, helpers in many cooperative breeders invest in pre-dispersal extra-territorial prospecting forays [e.g. 4, 5, 16, 22-24] whose conduct may trade-off against investments in natal cooperation [4, 5]. Such a trade-off between cooperation and activities that promote dispersal may effectively increase the direct fitness *costs* of cooperation for the more dispersive sex, given the greater fitness costs entailed in compromised dispersal for the more dispersive sex [5].

The Dispersal hypothesis therefore predicts that, in species in which both sexes delay dispersal and help in the natal group, individuals of the less dispersive sex should contribute to natal helping at higher rates. This prediction is distinct from the necessarily tight association between dispersal and helping in cooperative breeders in which only one sex ever delays dispersal and so only one sex *can* help within the natal group (a fairly common scenario in the cooperative birds [2]). While studies of individual species have revealed patterns consistent with the key prediction of the Dispersal hypothesis [e.g. 3, 5, 25-27], to what extent it can explain sex differences in helper effort within the natal group across cooperative taxa remains unclear. A recent comparative study has revealed that, among 15 cooperatively-breeding bird species in which both sexes help, a species’ sex difference in the probability of breeding within the natal group (as a subordinate or following inheritance of the dominant breeding position) predicts its sex bias in helper effort [6]; species with a more female-biased probability of natal breeding show more female-biased helper contributions. This finding holds promise for the Dispersal hypothesis, as sex biases in the probability of natal breeding likely arise at least in part via sex differences in dispersal. But whether dispersal patterns *per se* are driving this pattern is unclear, as sex biases in the probability of natal breeding will also be influenced by sex biases in reproductive skew within groups and rates of breeder turnover. Moreover, without restricting attention specifically to sex biases in helping quantified within the natal group (or while controlling for effects of variation in helper relatedness to recipients), any tendency for the more dispersive sex to help less could be attributable instead to a role for kin selection, given the higher likelihood that individuals of the more dispersive sex are immigrants, unrelated to within-group recipients. To establish whether the Dispersal hypothesis can indeed explain sex differences in helper contributions within the natal group, a dedicated comparative test of its key prediction is required.

Support for the Dispersal hypothesis would ideally stem from comparative support for its key prediction, coupled with evidence that such support cannot be readily attributed instead to confounding effects of other mechanisms that have been hypothesized to drive sex differences in helper effort in this context. Several other hypotheses have been proposed for the evolution of sex differences in helper effort (see [2] for a review, and the Discussion for wider coverage). For example, the ‘Heterogamety hypothesis’ [15] recognizes that while sons and daughters will be symmetrically related to recipients in their natal groups from the perspective of autosomal genes, the same need not be true for genes sited on the sex chromosomes. This observation leads to the prediction of greater helper effort in the homogametic sex; females in mammals and males in birds. Frequent observations of female-biased natal cooperation in mammals [e.g. 3, 25] and male-biased natal cooperation in birds [e.g. 26, 27] are therefore consistent with this hypothesis, while sex-reversals of these taxonomic norms give cause to question its central importance [e.g. 5]. The ‘Parental skills hypothesis’ proposes that the sex that contributes more to parental care might gain greater downstream direct fitness benefits from initially learning parenting skills via investment in helping [1]. While this is conceivable, the rarity of compelling evidence that helping actually does increase parental skills [2, 8, 28, 29] leaves the importance of this mechanism unclear. The ‘Paternity uncertainty hypothesis’ proposes that paternity uncertainty may favor greater investment in helping by males, by devaluing the expected fitness returns from breeding relative to helping to a greater extent in males than females [2, 30]. This hypothesis cannot therefore readily explain the evolution of female-biased cooperation [e.g. 3, 5, 25].

Here we use phylogenetic comparative methods to test the Dispersal hypothesis for the evolution of sex differences in cooperation, by assessing its ability to explain the patterns of sex differences in helper contributions to natal helping across cooperatively breeding mammals and birds. First, we use standard and phylogenetically controlled analyses to test the key prediction of the Dispersal hypothesis: among the cooperative breeders in which both sexes help within the natal group, a species’ sex difference in dispersal should predict its sex difference in contributions to natal cooperation, with helpers of the less dispersive sex contributing at higher rates to helping while within the natal group. Second, we assess whether the Dispersal hypothesis explains the cross-taxa patterns of sex differences in natal helping within the focal data set more effectively than the three other hypotheses outlined above: the Heterogamety, Parental skills and Paternity uncertainty hypotheses (in the Discussion we extend our attention to other hypotheses that are not yet tractable to test in a comparative context). Note that we are not seeking here to formally test the potential for the Heterogamety, Parental skills and Paternity uncertainty hypotheses to explain sex differences in cooperation more broadly, as that objective might be more effectively addressed with other (potentially larger) data sets ill-suited to testing the Dispersal hypothesis. Instead, our goal is to test the Dispersal hypothesis (which requires data to have been collected in specific contexts; see below), and to then establish whether the Dispersal hypothesis outperforms these alternative hypotheses in this context.

To test the Dispersal hypothesis, we focus our attention on species in which offspring of both sexes are known to help with offspring care while still residing within their natal group, and we seek to explain the direction of any sex difference in their rate of contributions to cooperative care in this context. We focused on helper contributions within the natal group as the rationale of the Dispersal hypothesis applies specifically to this context (see above) and because helpers of a given age within their natal group should not differ in their mean relatedness to recipients (reducing the likelihood that any sex difference in helper contributions is attributable instead to a role for kin selection [10]). This approach entailed restricting attention to the outcomes of studies that had statistically analyzed sex differences in the helper contributions (i) *solely* of individuals residing within their natal groups, or (ii) *also* including individuals with more diverse relationships to recipients while statistically controlling for effects of variation in helper relatedness to recipients (thereby rendering it unlikely that their findings were confounded by sex differences in mean relatedness to recipients; see Table S1 for details). We did not include species in which only one sex delays dispersal and so only one sex is *available* for natal helping, as the sex difference in dispersal necessarily predicts the sex difference in the occurrence of natal helping in such species [2, 8], and does so in the direction predicted by the Dispersal hypothesis. The inclusion of such species would therefore have falsely inflated any apparent support for the Dispersal hypothesis. This approach resulted in a sample size of 27 cooperatively breeding species (18 bird species and 9 mammal species) with which to test the Dispersal hypothesis. Full details of the species included and the source studies from which the relevant focal traits were collected can be found in Tables S1 and S2, along with relevant species-specific notes (see also Methods for full details of our approach to trait classification).

## METHODS

### Collating the Comparative Data Set

To collate the necessary data, we first collated an inclusive list of cooperatively breeding mammal and bird species from relevant reviews, comparative studies and books [9, 11, 31–34], before reducing this list via targeted species-by-species literature searches, to species in which (i) both sexes are known to help to rear the offspring of others within their natal group (or family/clan, in colonial species in which ‘groups’ can comprise multiple families/clans; e.g. sociable weavers, *Philataerus socius,* and white-fronted bee-eaters, *Merops bullockoides;* [35, 36]) *and* (ii) for which the sex bias in helper contributions to individual breeding attempts within the natal group (or family/clan) had been statistically investigated (or available data allowed its statistical investigation; see Table S1). This entailed restricting attention to source studies that had statistically analyzed sex differences in the helper contributions (i) solely of individuals residing within their natal groups, or (ii) also including individuals with more diverse relationships to recipients while statistically controlling for effects of variation in helper relatedness to recipients (thereby rendering it unlikely that their findings were confounded by sex differences in mean relatedness to recipients; see Table S1 for details). In some cases, it was necessary to seek confirmation from the authors of the source studies that the original analyses did meet these criteria, and in some cases the authors were able to provide revised analyses that did meet these criteria (see Tables S1 for full details; we greatly appreciate their assistance; see acknowledgements). For the species with analyses that met these criteria, the sex difference in helper contributions within the natal group had always been analysed without excluding members of the focal helper class that were not observed to contribute. As such, the source analyses should not have underestimated any sex differences in helper contributions that arise in part via a subset of individuals contributing nothing. For each species we restricted our attention to the outcomes of analyses of helping in contexts in which all available evidence suggested that the focal ‘helpers’ did not have young within the brood or litter that they were feeding (i.e. that they were indeed engaged in alloparental *helping* behaviour rather than parental care). For example, while our data set contains several species that are known to engage in joint-nesting / plural breeding (e.g. Seychelles warbler, *Acrocephalus sechellensis,* and acorn woodpecker, *Melanerpes formicivorus*), for such species we used the outcomes of analyses of the sex difference in helping among non-breeding helpers (see Table S1). So-called ‘failed-breeder cooperators’ (species in which helpers are typically individuals whose independent breeding attempts elsewhere have failed), were only included in our study if the sex difference in natal helping had been characterized at an age prior to the helpers dispersing away from the natal group or colony to attempt independent reproduction (as was the case for Western bluebirds, *Sialia Mexicana*, and white-fronted bee-eaters for example [35, 37]; Table S1), given the potential for complications to arise from any sex-specific dispersal already having occurred prior to the measurement of helping.

This approach yielded a data set of 27 cooperatively breeding species in which offspring of both sexes help while still resident in their natal groups (9 mammal and 18 bird species), and for which source studies had statistically tested for a sex difference in helper contributions within the natal group (see Tables S1 & S2 for full details of the relevant traits for these species, species-specific notes, and references to the source studies; we regret that we cannot cite all of the source studies within the main paper too, due to restrictions on reference numbers). These 27 species were used as the focal data set for our study. For a further nine species in which offspring of both sexes are known to help within their natal groups, source studies had statistically analysed the sex difference in helper contributions, but *without* restricting attention to helpers within their natal groups and without controlling for variation in helper relatedness to recipients, leaving their outcomes ill-suited to testing the Dispersal hypothesis (as any relationship between sex differences in dispersal and helper contributions in these species could be confounded by sex differences in the mean relatedness of helpers to recipients within the helping data sets analysed; see rationale above). As such, these nine species were not included in our analyses and play no further role in our study. In case of interest (or utility for other studies), we do still report these species and their sex differences in dispersal and helping (which may not reflect their patterns of *natal* helping) within our Supplementary Tables S1 & S2 (the grey-highlighted species at the end of each table), where one can see that their sex differences in dispersal and helping nevertheless echo the patterns within our analysed data set of 27 species in being broadly consistent with the predictions of the Dispersal hypothesis (e.g. see footnote 7 in Tables S1).

For all 27 focal species, we collated information from original source studies (see Tables S1 and S2) on the (i) sex difference in dispersal from the natal group (or family), categorized according to whether dispersal (assessed via multiple traits; see below) was significantly male biased (MB), showed no significant sex bias (NSB), or was significantly female biased (FB), and (ii) sex difference in helper contributions within the natal group (specifically, the rate or incidence of offspring care provision by natal helpers [see below]; again, MB, NSB, or FB). The sex differences in both traits were scored in this categorical way as effect sizes or the data required to calculate them were not always available and could vary due to among-study variation in the focal dispersal and cooperation traits assessed (see below) or the covariates fitted in the original analyses. Wherever information was available for the sex difference in helper contributions to more than one form of offspring care (e.g. both incubating and nestling feeding), a species was classified as showing a sex difference in helper contributions if a significant sex difference was apparent in one or more forms of helper care (for none of the 27 species did two forms of care by helpers exhibit significant sex biases in opposite directions; see Table S1 for details). Sex differences in dispersal were scored by examining statistical analyses of sex differences in several dispersal-related traits: incidence of dispersal from the natal territory, age at first dispersal, dispersal distance and population genetic structure as it relates to dispersal (see Table S2 for details). A species was classified as showing a sex difference in dispersal if a significant sex difference was apparent in one or more of these metrics in one or more studies (as sex biases in *any* of these metrics could impact the net direct fitness payoff from natal cooperation via the mechanisms envisaged in the Dispersal hypothesis, which span the impacts of dispersal patterns on both the benefits and costs of helping; see Introduction). Again, for no species did two of these dispersal traits show significant sex biases in opposite directions.

To then allow comparisons of the explanatory power of the Dispersal, Heterogamety, Parental skills and Paternity uncertainty hypotheses in this specific context, we also collated information for the focal 27 species on (i) whether the species was a bird (females heterogametic) or a mammal (males heterogametic), (ii) the sex bias in parental contributions to the same form of offspring care as that used for the assessment of sex biases in helper contributions (again, MB, NSB, or FB), and (iii) the incidence of both extra-pair and extra-group paternity (see Tables S1 & S2 for details). When collating the paternity data, we recorded both the average proportion of offspring not sired by the dominant breeding male within the group (‘extra-pair paternity’) and the average proportion of offspring not sired by any male within the group (‘extra-group paternity’). We then assessed the explanatory power of the Paternity uncertainty hypothesis using both data forms, as (i) this hypothesis [2, 30] relates to the paternity certainty of the breeding male (rendering extra-pair paternity a relevant currency), but (ii) assumes that any paternity lost is lost to an unrelated male, leaving extra-group paternity a potentially more appropriate currency because a proportion of extra-pair paternity could be won by within-group males related to the dominant male (see [11] for a similar argument).

### Phylogenetic logistic regression analysis

We initially assessed support for the Dispersal hypothesis with a simple contingency table analysis (using Fisher’s exact tests, given the modest sample sizes), which allowed us to assess the association between the trinomial predictor (sex bias in dispersal: MB, NSB or FB) and response (sex bias in helping: MB, NSB or FB) variables, but without controlling for phylogenetic effects. We then used phylogenetic logistic regressions to assess the key prediction of the Dispersal hypothesis while controlling for the phylogenetic non-independence of species [38], by recategorizing our trinomial response variable (sex bias in helping; MB, NSB or FB) into two binary response variables. To do this we created two new sex-bias-in-helping variables (FB helping versus Other [MB & NSB combined], and MB helping versus Other [FB & NSB combined]) and used these to test whether the sex bias in dispersal (MB, NSB or FB) predicted the sex bias in natal helping using both binomial helping classifications (i.e. FB help vs Other, and MB help vs other). The phylogenetic logistic regressions were run using the phyloglm function in the R package phylolm [39], and accounted for phylogenetic non-independence by incorporating a variance-covariance matrix based on the structure of the phylogenetic tree (see below for details). Standard (non-phylogenetic) logistic regression models were also run. Following Ives & Garland (2010) we ran bootstrap analyses with 100 replicates in the phylogenetic logistic regression to derive means and 95% confidence intervals for the parameters. The relative strength of support for competing models (reflecting the competing hypotheses outlined above) was assessed using an information theoretic approach, in which AICc-based assessments of model fit were determined for all models. Following Burnham & Anderson [40] a ΔAICc > 2 between pairs of competing models was considered to reflect stronger statistical support for the model with the lower AICc score.

First, we assessed the strength of support for the Dispersal hypothesis (i.e. a model fitting the sex bias in dispersal [MB, NSB or FB] as the sole predictor other than phylogeny) relative to a null (phylogeny only) model, using the full data set of 27 species. Second, we use the same phylogenetically-controlled approach to compete the Dispersal hypothesis model against models capturing the three other hypotheses outlined above. The explanatory power of the Dispersal hypothesis model was compared to that of (i) a Heterogamety hypothesis model (i.e. fitting whether the species was a bird [females heterogametic] or a mammal [males heterogametic] as the sole predictor other than phylogeny) using the full data set of 27 species, (ii) a Parental skills hypothesis model (i.e. fitting the sex bias in parental contributions to the same form of care as assessed for helping [MB, NSB or FB] as the sole predictor other than phylogeny) using the reduced data set of 21 species for which the necessary parental contributions data was available (see Tables S2), and (iii) a Paternity uncertainty hypothesis model (i.e. fitting the incidence of extra-pair or extra-group paternity as the sole predictor other than phylogeny) using a reduced data set of 18 species. These 18 species were those for which the necessary paternity data was available, following the exclusion of five joint-nesting species (as while helping in the joint-nesting species included was assessed among non-breeding helpers [see above], joint nesting systems could generate maternity uncertainty, voiding a key assumption of the Paternity uncertainty hypothesis [30]).

### Phylogenetic trees and assessment of alternative phylogenetic assumptions

Information on the phylogenetic relationships between the species included in this study was taken from composite phylogenetic trees (“Supertrees”) of mammals [41] and birds [42]. We also checked an alternative source for the bird phylogenies [43], which showed the same relative relationships between the species examined in this study. Eight of the 27 species (6 birds, 2 mammals) considered were not contained in these trees (see Table S3). For six of these cases the missing species could be assigned the phylogenetic position of a suitable closely-related species (e.g. a sister species or congener; see Table S3). In the remaining two cases species were assigned to a new branch in a position in the tree based on their standard taxonomic classification (Table S3). To enable mammals and birds to be included in the same analyses, the mammalian and avian trees were joined together such that their root nodes were joined by branches to a new root node representing their common ancestor.

In creating these trees and running the phylogenetic comparative analyses, assumptions about the branch lengths of the phylogenetic tree needed to be made. In this context, branch lengths represent the degree of divergence between species either in time or in the data type used to infer the phylogeny (typically genetic data). In the mammal supertree the supplied branch lengths were in units of time, but branch lengths were not supplied in the bird supertree. In order to join these trees and include the additional taxa mentioned above we therefore initially set all branch lengths in the combined mammal and bird tree to be equal (i.e. all branch lengths take a value of one). This has the effect of assuming a short evolutionary distance between bird and mammal lineages. Given that birds and mammals last shared a common ancestor more than 300 million years ago [44], we explored the effect on our analyses of extending these two branch lengths (from both mammals and birds back to their common ancestor). To do this, we created three new trees with branch lengths leading from the root (common ancestor) to the birds and the mammals that were 5, 10 or 100 times as long as the other branches (Figure S1). Rerunning our phylogenetic analysis using these alternative trees confirmed that our findings are robust to variation in these branch lengths to the common ancestor (Table S4). All analyses presented in the main text are therefore based on the simplest assumption of equal branch lengths (though further tests were also conducted to verify that our findings are robust to the nature of this assumption; see below).

Our analyses use the Ives & Garland [45] approach to estimating the phylogenetic logistic regression. This method estimates the degree to which phylogenetic effects need to be controlled for, by estimating a “phylogenetic signal” parameter *a*, which increases with increasing magnitude of the phylogenetic correlations among species. The potential range for the parameter *a* spans zero, with more positive values indicating stronger phylogenetic signal and more negative values indicating weaker phylogenetic signal, with values below −4 considered to indicate negligible phylogenetic signal [45]. We explored the effects on our findings of making different phylogenetic assumptions and using different analytical approaches. Specifically, we investigated the effects of (i) further variation in the assumptions regarding branch lengths (we used Grafen’s method for arbitrarily creating ultrametric trees based on the tree structure [46], exploring the effects of 3 values of the Rho scaling parameter which affects the relative length of branches based on their closeness to the root of the tree; Figure S2), (ii) explicitly assuming that there is no phylogenetic signal in the data (implemented in two ways: first by scaling the branch lengths by Pagel’s λ with λ=0, and second by conducting standard, non-phylogenetic logistic regression), and (iii) using Maximum Penalized Likelihood estimation (again with all branch lengths set to one; see above) instead of the Ives & Garland [45] method. Using these alternative approaches had no effect on the relative support that the analyses revealed for the competing hypotheses, perhaps principally because our analyses suggest that there was only weak phylogenetic signal in the data set (see Results). The AICc values for all of the competing models are reported for each one of these different approaches within Tables S5 & S6.

## RESULTS

### Testing the Dispersal hypothesis

To test the Dispersal hypothesis, we compiled a data set of 27 species in which non-breeding helpers of both sexes help to rear offspring within their natal group, and for which source studies had statistically tested for the existence of sex differences in both (i) dispersal (significant Male-bias [MB], no significant sex bias [NSB], significant Female-bias [FB]) and (ii) helper contributions within the natal group (MB, NSB or FB). The 27 species were taxonomically diverse (Figure 1), comprising 18 bird and 9 mammal species, representing 24 genera, 19 families and 6 orders. Notably, in all 13 species that showed significant sex biases in both dispersal and natal helping effort, species with male-biased dispersal showed female-biased natal helper contributions, while species with female-biased dispersal showed male-biased natal helper contributions (Figures 1 and 2a), just as predicted by the Dispersal hypothesis. Not a single species showed the opposite sex bias in natal helping to that which would be predicted by the Dispersal hypothesis. Contingency table analyses (i.e. lacking phylogenetic control) confirmed that a species’ sex bias in dispersal (MB, NSB or FB) significantly predicted its sex bias in natal helping (MB, NSB or FB; n = 27 species, Fisher’s 3×3 exact test p < 0.001; Figure 2a). As predicted by the Dispersal hypothesis, species with male-biased dispersal were significantly more likely to show female-biased natal helping (7 of 9 species) than species with female-biased dispersal (0 of 14 species; Fisher’s 2×2 exact test p < 0.001), while species with female-biased dispersal were significantly more likely to show male-biased natal helping (6 of 14 species) than species with male-biased dispersal (0 of 9; Fisher’s 2×2 exact test p = 0.048).

**Figure 1.**
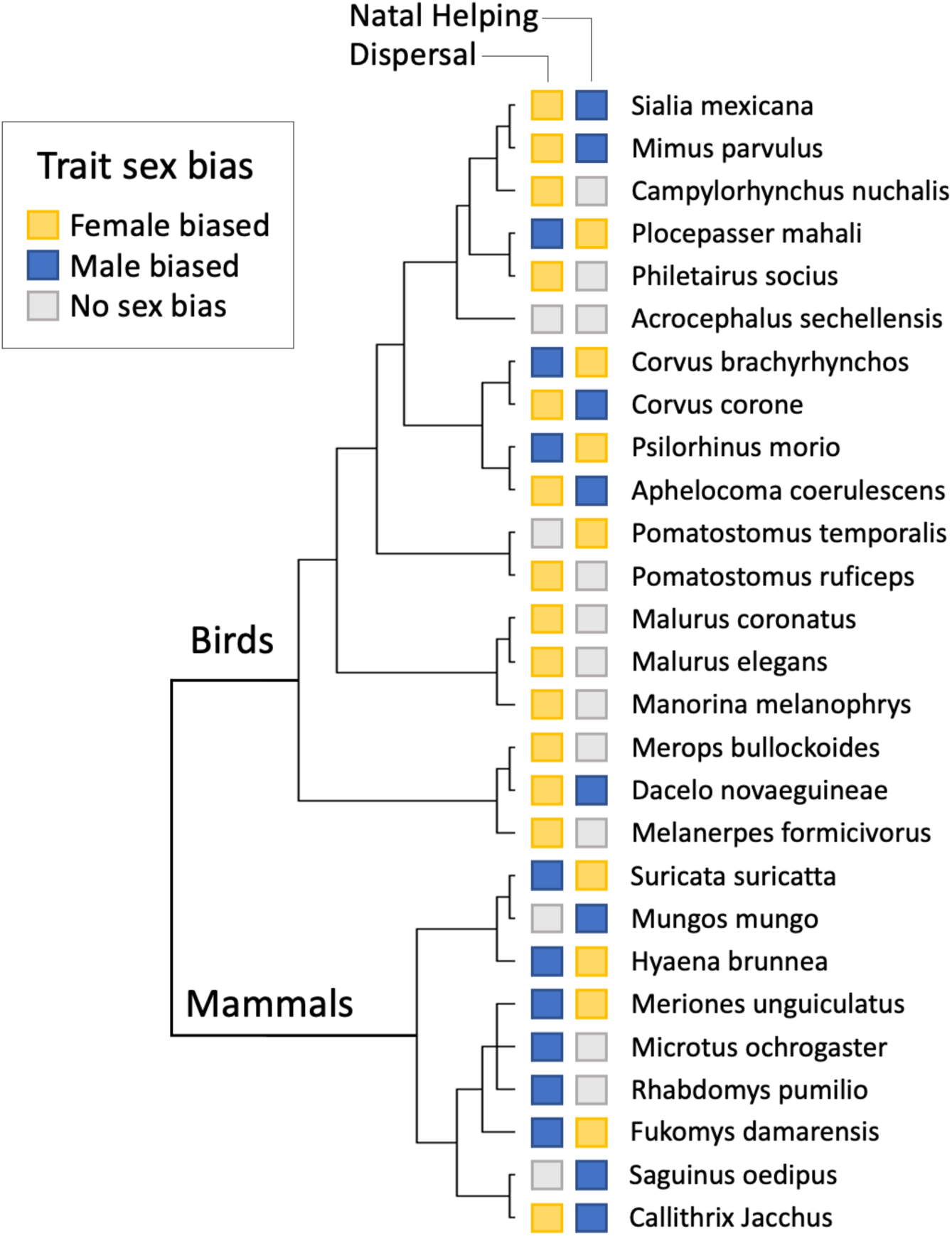
The phylogenetic distribution of the 27 focal species in the analyses within the main paper, representing 24 genera, 19 families and 6 orders. The shaded squares present our codings for the species’ sex biases in Dispersal (left) and Natal Helping (right), which reflect whether the source studies found a statistically significant sex bias in the focal trait and the direction of any significant sex bias detected (see methods and Tables S1 & S2 for details). Notably, every species that shows significant sex differences in both traits shows the sex difference in natal helping that would be predicted by the Dispersal hypothesis on the basis of its sex difference in dispersal (see also Figure 2a). All other species show no significant sex bias in either dispersal or natal helping. Not a single species shows the opposite sex bias in natal helping to that which would be predicted by the Dispersal hypothesis. The tree captures the structure of the phylogenetic relationships in the tree but not the branch lengths tested (for which a range of different assumptions were tested; see Methods and Tables S5 and S6).

**Figure 2.**
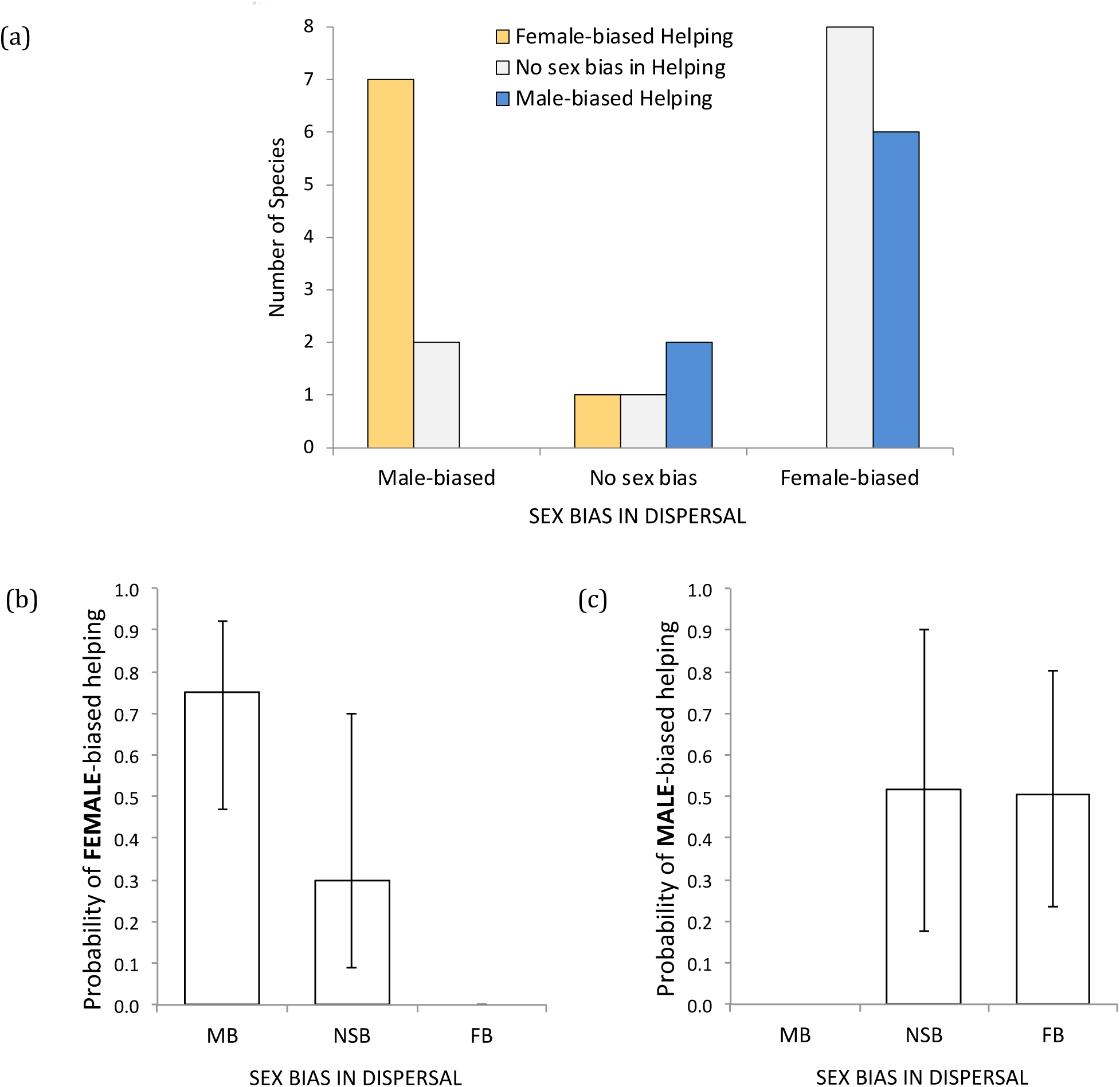
The Dispersal hypothesis predicts that, among helpers within their natal group, the more dispersive sex should contribute less to helping. Our comparative analyses of 27 species of cooperative birds and mammals support for this prediction. (**a**) Female-biased helping was strongly associated with male-biased dispersal (and never occurred alongside female-biased dispersal), while male-biased helping was strongly associated with female-biased dispersal (and never occurred alongside male-biased dispersal). Accordingly, in our phylogenetic comparative analyses, the sex bias in dispersal significantly predicted whether natal helper effort was (**b**) significantly *Female*-biased or not, and (**c**) significantly *Male*-biased or not (see results). In (b) and (c) the bars present predicted means (± 95% confidence intervals) while controlling for phylogenetic effects. MB = (significantly) Male-biased, NSB = No significant sex bias, FB = (significantly) Female-biased.

Phylogenetic comparative analysis confirmed that these relationships hold when controlling for phylogenetic effects: the sex bias in dispersal (MB, NSB, or FB) predicted both the probability of female-biased natal helping (Figure 2b; ΔAICc = −13.40 relative to the phylogeny-only model; R^2^ = 0.61, n = 27 species, a = −2.41 indicating weak phylogenetic signal) and the probability of male-biased natal helping (Figure 2c; ΔAICc = −3.53 relative to the phylogeny-only model; R^2^ = 0.26, n = 27 species, a = −2.17 indicating weak phylogenetic signal). Inspection of the estimates and 95% confidence intervals for the predicted means (Figures 2b and 2c) and the pair-wise contrasts among dispersal classes (Table 1) confirms support for the Dispersal hypothesis. First, in the model of the probability of female-biased helping (Table 1 upper half; Figure 2b), species with male-biased dispersal were significantly more likely to show female-biased helping than species with female-biased dispersal (while species with NSB dispersal showed an intermediate probability of female-biased helping). Second, in the model of the probability of male-biased helping (Table 1 lower half; Figure 2c), species with female-biased dispersal were significantly more likely to show male-biased helping than species with male-biased dispersal (while species with no significant sex-bias in dispersal showed a similar probability of male-biased helping to those with female-biased dispersal). These findings were robust to (i) using different assumptions regarding phylogenetic branch lengths (Tables S4-S6), (ii) using standard logistic regression without controlling for phylogeny (Tables S5 & S6), and (iii) using the maximum penalized likelihood estimation method for phylogenetic logistic regression in place of the Ives & Garland [45] method (Tables S5 & S6).

**Table 1.** Effect size estimates from the phylogenetic logistic regressions investigating whether sex biases in dispersal predict sex biases in helping within the natal group.

| Response Variable | Parameters | Estimated effect size | Boot-strapped 95% CI |
| --- | --- | --- | --- |
| **Probability of Female-biased Helping**<br>(vs Other) | *Intercept* | 2.06 | -1.70, 5.83 |
|  | *Sex bias in Dispersal* |  |  |
|  | Male-biased (reference) | 0.00 | - |
|  | No Sex Bias | -2.88 | -7.11, 1.34 |
|  | Female-biased Dispersal | -18.80* | -20.54, -17.06 |
| **Probability of Male-biased Helping**<br>(vs Other) | *Intercept* | -19.85 | -21.43, -18.28 |
|  | *Sex bias in Dispersal* |  |  |
|  | Male-biased (reference) | 0.00 | - |
|  | No Sex Bias | 20.34* | 18.43, 22.25 |
|  | Female-biased Dispersal | 20.04* | 18.50, 21.59 |

The sex bias in dispersal significantly predicted both (i) the probability that helper contributions within the natal group were significantly female-biased (upper half of table; ΔAICc = −13.40 relative to the phylogeny-only model; n = 27 species; Figure 2b; Table S5) and (ii) the probability that helper contributions within the natal group were significantly male-biased (lower half of table; ΔAICc = −3.53 relative to the phylogeny-only model; n = 27 species; see also Figure 2c; Table S6). The effect size estimates for the contrasts between a given dispersal factor level (either female-biased dispersal or no significant sex bias in dispersal) and the reference level (male-biased dispersal) are shown here, along with their boot-strapped 95% confidence intervals. The effect sizes for factor levels of the sex bias in dispersal predictor whose estimates differ significantly from that for male-biased dispersal (the reference level) are highlighted *. These models used the Ives & Garland [45] method with all branch lengths set = 1 (see Methods).

### Testing the Dispersal Hypothesis against Alternative Explanations

Comparisons of alternative (phylogenetically controlled) models for explaining the incidence of *female*-biased helping revealed substantially stronger support for the Dispersal hypothesis than the Heterogamety, Parental skills and Paternity uncertainty hypotheses. Heterogamety hypothesis: sex bias in dispersal was a stronger predictor of female-biased helping than whether the focal species was a bird (females heterogametic) or mammal (males heterogametic; ΔAICc = −14.58; n = 27 species with data for both predictors; Table S5 upper third). The Heterogamety hypothesis model (i.e. allowing for an effect of whether the species was a bird or mammal) explained the data no more effectively than the null (phylogeny only) model (ΔAICc = +1.18; n = 27; Table S5 upper third). As this null (phylogeny only) model may itself account for effects of any contrast between birds and mammals via the phylogeny, we note that the Heterogamety hypothesis model also explained the data no more effectively than the null model when phylogenetic effects were not controlled (e.g. setting lambda = 0, ΔAICc = +1.18; or using standard logistic regression, ΔAICc = +0.89; Table S5 upper third). Parental skills hypothesis: sex bias in dispersal was also a stronger predictor of female-biased helping than sex bias in parental care (ΔAICc = −12.73; n = 21 species with data for both predictors; Table S5 middle third). The Parental skills hypothesis model (i.e. allowing for an effect of the species’ sex bias in parental care) explained the data no more effectively than the null, phylogeny only, model (ΔAICc = +4.69, n = 21 species; Table S5 middle third). Paternity uncertainty hypothesis: sex bias in dispersal was also a stronger predictor of female-biased helping than the incidence of paternity loss (ΔAICc relative to extra-*pair* paternity predictor = −12.93; ΔAICc relative to extra-*group* paternity predictor = −12.54; n = 18 species with paternity data; Table S5 lower third). The Paternity uncertainty model (i.e. allowing for an effect of the extent of paternity loss) did not explain the data significantly more effectively than the null, phylogeny only, model (ΔAICc for extra-*pair* paternity = −0.95; ΔAICc for extra-*group* paternity = −1.34; n = 18 species; Table S5 lower third). The patterns in the raw data (Figure S2b) suggest that any weak association that there is between paternity uncertainty and sex biases in helping actually runs counter to the predictions of the Paternity uncertainty hypothesis (species with female-biased helping if anything show the highest incidence of extra-pair/group paternity; Figure S3c). The outcomes of these model comparisons were robust to using a range of different phylogenetic assumptions (Table S5). The raw data associations between the Heterogamety, Parental skills and

Paternity uncertainty hypothesis predictors and species’ sex biases in helping are presented in Figure S2. Comparisons of alternative (phylogenetically controlled) models for explaining the incidence of *male*-biased helping revealed that the Dispersal hypothesis model outperformed both the null (phylogeny only) model (ΔAICc = −3.53; n = 27 species; Table S6 upper third) and the Heterogamety hypothesis model (ΔAICc = −6.07; n = 27 species; Table S6 upper third) when using the full data set of 27 species. Once again, the Heterogamety hypothesis model explained the data no more effectively than the null model, whether controlling for phylogenetic effects or not (ΔAICc = +2.18 to +2.55 depending on the method; n = 27; Table S6 upper third). Using the reduced data set available for testing the Parental skills hypothesis (n = 21 species), neither the Dispersal nor Parental skills hypothesis models consistently outperformed (ΔAICc < −2) the null, phylogeny only, model across all sets of phylogenetic assumptions (Table S6 middle third). While the Parental skills hypothesis did outperform the null model in two phylogenetic scenarios (for Rho = 3 and Standard logistical regression [i.e. without phylogenetic control]; Table S6 middle third) it did not outperform the Dispersal hypothesis model in any of the scenarios tested. When using the reduced data set available for testing the Paternity uncertainty hypothesis (n = 18 species), neither the Paternity uncertainty hypothesis nor the Dispersal hypothesis predicted the incidence of male-biased helping significantly more effectively than the null (phylogeny only) model (ΔAICc > −1 for all comparisons to the null model under all phylogenetic assumption sets tested; Table S6 lower third).

## DISCUSSION

Our comparative analyses sought to test the Dispersal hypothesis for the evolution of sex differences in cooperation, which proposes that the more dispersive sex should contribute less to natal helping in any given breeding attempt as it may stand to gain a lower net direct fitness payoff from natal helping (because dispersal can impact the downstream direct benefits and/or costs of natal cooperation; see Introduction). Our analyses therefore focused on species in which non-breeding helpers of both sexes help to rear offspring within their natal groups, and investigated whether, across such taxa, sex differences in dispersal predict sex differences in helper contributions within the natal group. Notably, in every species that showed significant sex differences in both dispersal and natal helping, the sex bias in dispersal predicted that in natal helping in the direction predicted by the Dispersal hypothesis. Accordingly, our phylogenetic comparative analyses revealed support for the Dispersal hypothesis. Species’ sex biases in dispersal significantly predicted their sex biases in natal helping, when modelling both the probability of female-biased helping (Figure 2b) and the probability of male-biased helping (Figure 2c). As predicted, relative to species with male-biased dispersal, species with female-biased dispersal were significantly *more* likely to show male-biased contributions to natal helping (Figure 2c; Table 1 lower half) and significantly *less* likely to show female-biased contributions to natal helping (Figure 2b; Table 1 upper half). This association is distinct from the necessarily perfect association between sex biases in dispersal and sex biases in natal cooperation in those cooperative breeders in which only one sex delays dispersal and so only that sex is available for natal helping (as such species were not included in the analysis). Support for the Dispersal hypothesis was also robust to varying the phylogenetic assumptions within our models (see methods, Figures S1 & S2 and Tables S4-6). Below, we consider potential alternative explanations for our findings, highlight that the mechanisms by which sex differences in dispersal might drive sex differences in natal cooperation demand closer attention, and consider the wider implications of these findings for our understanding of the evolution of cooperation.

Having found support for the Dispersal hypothesis, we tested whether such support could be attributed instead to confounding effects of the mechanisms envisaged in the Heterogamety, Parental skills and Paternity uncertainty hypotheses. Our analyses suggest that this is unlikely, as (i) none of these other hypotheses consistently outperformed a null model in any of our analyses, and (ii) the Dispersal hypothesis significantly outperformed all of these hypotheses, as well as the null model, either in the model of female-biased helping, the model of male-biased helping, or both. That the Dispersal hypothesis model outperformed the bird/mammal contrast captured in the Heterogamety model is notable, as sex biases in dispersal could have been confounded by a bird/mammal contrast, given that birds and mammals often show female- and male-biased dispersal respectively [47]. Indeed, species that show rare examples of male-biased dispersal among birds (e.g. brown jays, *Psilorhinus morio* [5, 48] and American crows, *Corvus brachyrhynchos* [49]) illustrate the predictive power of the Dispersal hypothesis, as these species are also unusual among cooperative birds in showing female-biased contributions to natal helping [5, 49]. When using the restricted data sets available for competing the Dispersal hypothesis against the Parental skills (n = 21 species) and Paternity uncertainty (n = 18 species) hypotheses, the Dispersal hypothesis outperformed both (and the null model) when modelling the probability of female-biased helping, but none of these hypotheses consistently outperformed the null model when modelling male-biased helping. This latter ambiguity could be due to the restricted sample sizes available for these comparisons, coupled with patterns of dispersal appearing to be a slightly stronger predictor of female-biased helping than male-biased helping in our data set (a pattern echoed in our analyses of the full data set). The overall lack of support for the Heterogamety, Parental skills and Paternity uncertainty hypotheses in our analyses suggests that they do not provide credible alternative explanations for our findings in support of the Dispersal hypothesis (the focal hypothesis under test here) and are unlikely to be the *primary* driver of sex differences in cooperation in the context studied here. Our findings do not give cause to rule out *any* role for the mechanisms envisaged in these alternative hypotheses though, as they could conceivably still have contributed to selection for sex differences in cooperation in this and other contexts (see Supplementary Material - Extended Discussion).

It has also been suggested that the sex that shows higher variance in lifetime reproductive success (hereafter termed ‘reproductive variance’) may invest more in helping early in life as their chance of ultimately securing direct fitness by becoming a breeder is lower ([1], see also [2]). This idea warrants formal modeling, however, as it seems conceivable that the sex with higher reproductive variance could also stand to gain more from *foregoing* helping, because a given consequent increase in competitive ability (e.g. if helping entails costs to growth and/or body condition; [50–52]) could presumably yield a greater downstream direct fitness return in the sex with higher reproductive variance. Either way, while the rarity of studies that have estimated sex differences in reproductive variance in cooperative breeders currently precludes comparative tests of this idea [53], the limited available evidence does not support a simple association between the sex biases in reproductive variance and either dispersal or natal helping effort in the types of species studied here (those in which both sexes delay dispersal and help in the natal group). Data for meerkats, *Suricata suricatta,* and Damaraland mole-rats, *Fukomys damarensis*, for example, appear consistent with the hypothesis, as reproductive variance is estimated to be higher in females than males in both species [53, 54] and females are also the less dispersive and more helpful sex [3, 25, 55]. However, females have also been estimated to show higher reproductive variance than males in superb starlings, *Lamprotornis superbus* [56], where males are the more philopatric sex and helper contributions appear to be male biased [57].

The observed association between sex biases in dispersal and natal cooperation also cannot be readily attributed instead to helpers being studied in contexts in which those of the more dispersive sex are on average less related to their recipients than those of the less dispersive sex (a scenario in which kin selection might account for our findings). This is because our analyses focused specifically on studies that had characterized sex differences in the contributions of helpers (i) within their natal groups (where male and female helpers should not differ in their mean autosomal relatedness to recipients) or (ii) while controlling for effects of variation in relatedness to recipients if helpers in more diverse relatedness contexts were also included. Another potential complication is that the less dispersive sex is likely to stay for longer on average in its natal group, and so the original source studies could have monitored natal helping on average at older ages in this sex than the more dispersive sex (leaving sex differences in dispersal confounded by sex differences in mean helper age at monitoring). However, we minimized the potential for such an age confound by drawing wherever possible on analyses of natal helping effort that had controlled for effects of helper age (either by restricting attention to a given age class or by statistically controlling for age effects; see Table S1). While this was not possible for four of the 27 species used (Table S1), follow-up analyses confirmed that removing these species from the data set left the outcomes of our model comparisons unchanged (see footnote 5 in Table S1). As such, our findings cannot be readily attributed instead to sex differences in mean age-at-monitoring within the original studies.

Our analyses provide support for the Dispersal hypothesis, which proposes that sex differences in natal cooperation evolve because the more dispersive sex stands to gain a lower net direct fitness payoff from natal helping, due to two potentially widespread mechanisms that could act in isolation or concert in any given species. The more dispersive sex may (i) stand to gain a lower direct fitness *benefit* from natal helping [3, 6] and/or (ii) experience a greater direct fitness *cost* of natal helping [4, 5]. The more dispersive sex could gain a lower direct *benefit* from natal helping wherever accruing a direct benefit is contingent in part upon continued philopatry. This could be the case either because helping improves a local public good (e.g. the size of the natal group or territory, from which residents may derive a direct benefit as long as they remain within the group, particularly if they breed there [3, 6, 16, 20, 21]) or because the benefits of helping are contingent in part upon continued interactions with prior recipients or observers of one’s help (e.g. a role for reciprocal altruism, the accrual of social prestige, or signaling one’s quality to mates [2, 8]). The more dispersive sex could also suffer a greater direct fitness *cost* of helping, because investment in helping may trade off against their simultaneous need to invest in activities that promote dispersal (such as extra-territorial prospecting, growth or the accrual of body reserves; [4, 5]). This latter mechanism, focused on direct costs of helping, could arguably apply more widely than the former (focused on direct benefits of helping), as it does not require that helping yields a downstream direct benefit whose magnitude is contingent upon remaining in the natal group. Our comparative findings do not allow us to discriminate between these two mechanisms. Similarly, while recent evidence that the sex bias in probability of natal breeding predicts the sex bias in helping across 15 cooperative bird species could reflect a role for sex differences in the direct *benefits* of helping (which could scale with the probability of natal breeding; see above) [6], such a pattern could also reflect a role for sex differences in the direct fitness *costs* of helping (because the sex that is less likely to breed within the natal group may invest more in activities that promote dispersal, at the expense of natal cooperation [4, 5]). This pattern is nevertheless consistent with the predictions of the Dispersal hypothesis, to the extent that the focal sex biases in probability of natal breeding (both as a subordinate and following dominance acquisition; their effects could not be teased apart [6]) arose via sex differences in dispersal rather than sex differences in reproductive skew and/or dominance tenure length. While the two general mechanisms by which sex differences in dispersal could drive sex differences in natal cooperation (i.e. via sex differences in the direct fitness benefits or costs of cooperation) are likely to prove difficult to tease apart, attempts to do so might now be usefully prioritized. Indeed, variation among species in the relevance of these two mechanisms could also help to explain deviations within our data set from the patterns predicted by the Dispersal hypothesis (see Supplementary Materials – Extended Discussion).

Our findings highlight a contemporary association between sex differences in dispersal and natal cooperation that is consistent with the scenario envisaged in the Dispersal hypothesis, in which past evolutionary changes in sex differences in dispersal drove changes in the sex difference in natal cooperation. However, it is conceivable that sex differences in natal helping have also (or instead) shaped the patterns of selection on sex differences in dispersal. For example, wherever trade-offs exist between investments in helping and dispersal-promoting activities [4, 5], sex differences in natal cooperation would also have the potential to drive sex differences in dispersal. While the co-evolution of sex differences in dispersal and cooperation does seem plausible, it seems unlikely that the association observed here *solely* reflects sex biases in cooperation driving sex biases in dispersal. Such a scenario would require the evolution of significant sex biases in natal helping via some alternative mechanism (other than those envisaged in the Dispersal hypothesis) that then drove evolutionary changes in the sex biases in dispersal, bringing the species’ sex biases in both traits in line with the predictions of the Dispersal hypothesis. It is notable then that we currently lack compelling support for such alternative mechanisms for the evolution of sex differences in cooperation (see above; e.g. the Heterogamety, Parental skills and Paternity uncertainty hypotheses). Moreover, most species in our data set have sex biases in dispersal that conform to their historical taxonomic norms (i.e. female-biased dispersal in birds and male-biased dispersal in mammals [47]), leaving it likely that the dispersal patterns of most of our focal species were in place prior to the evolution of helping in their clade. That said, our findings do highlight the need for closer attention to the possibility that sex differences in the payoff from natal cooperation influence the patterns of selection on dispersal [8, 58].

Together, our analyses provide novel support for the Dispersal hypothesis for the evolution of sex differences in cooperation. Our analyses necessarily focused on the subset of cooperative breeders in which both sexes help within their natal group, because the rationale of the Dispersal hypothesis only applies in this context. Nevertheless, the observed association between sex differences in dispersal and natal cooperation will of course extend to cooperative breeders in which only one sex delays dispersal, as in such species only members of the less dispersive sex are *available* to engage in natal cooperation. While there is naturally a need for caution in generalizing our findings beyond the subset of species for which the necessary data were available, our findings do highlight the potential for sex differences in dispersal to have played a widespread role in driving sex differences in cooperation across taxa (whether in isolation or acting in concert with other mechanisms). As our analyses focused on helping behaviour within the natal group, it remains an open question to what extent patterns of dispersal are also relevant to understanding sex differences in (i) other forms of cooperation (e.g. cooperative vigilance, foraging, construction and territorial defence [59–64]) and (ii) cooperation in other contexts (e.g. following dispersal from the natal group [14, 19, 59, 61, 65, 66]). While a wealth of evidence now supports the view that *indirect* fitness benefits have played a key role in the evolution of cooperation via kin selection [8, 11–13], our findings provide evidence suggestive of a wider role for differences in net *direct* fitness payoffs in shaping patterns of cooperation across species [6, 19, 66, 67]. Importantly though, our findings do not implicate a role specifically for sex differences in the direct fitness *benefits* of cooperation, as sex differences in dispersal and philopatry could also drive sex differences in cooperation by generating sex differences in the direct fitness *costs* of cooperation [4, 5]. A key focus for future work on individual model systems will therefore be to establish whether sex differences in dispersal shape patterns of cooperation via impacts on the sex-specific direct fitness benefits or costs of cooperation.

## ACKNOWLEDGEMENTS

We thank Mike Cant, Pablo Capilla-Lasheras, Jeremy Field, Xavier Harrison, Sarah Hodge, Shakti Lamba and Andy Russell for fruitful discussions and/or comments on the manuscript, and Elena Berg, Lyanne Brouwer, Steve Emlen, Anne Peters, Morne du Plessis, Andy Radford, Mandy Ridley, Dustin Rubenstein, Sheng-Feng Shen and Faye Thompson for responses to queries during data compilation. AJY was supported by a BBSRC David Phillips Research Fellowship (BB/H022716/1).

