## Supplementary Materials for "Sex differences in dispersal predict sex differences in helping across cooperative birds and mammals"

*Fenner et al – Sex differences in dispersal predict sex differences in cooperation in cooperatively breeding birds and mammals*

**Table S1 – Data for sex biases in helper contributions to cooperation**

The first 27 species in the table were used in the analyses presented in the main paper. They constitute the cooperatively breeding bird and mammal species in which both sexes help as non-breeders within their natal group, for which source studies had statistically tested for sex differences in both dispersal (Table S2) and helper contributions within the natal group (or, where helpers of other classes were also included, while controlling for effects of variation in relatedness to recipients). The last 9 species in the table (highlighted in grey) were *not* used in the analyses in the main paper, as they meet the same criteria *except* the source studies' statistical analyses of the sex difference in helper contributions were *not* restricted to helpers within their natal group and did *not* control for variation in helper relatedness to recipients, leaving their outcomes ill-suited to testing the Dispersal hypothesis (as sex differences in dispersal in these species could be confounded by sex differences in mean helper relatedness to recipients within the source analyses; see Methods in the main paper). We include these species here to clarify why they weren't included in our analyses and to show that their cooperation (this table) and dispersal (Tables S2) sex differences are also consistent with the predictions of the Dispersal hypothesis (see table footnote 7). 'Female-biased' and 'Male-biased' refer to statistically significant sex differences. 'NSB' refers to no significant sex difference. See table footnotes and Methods for further details.

| Species | Sex-bias in Helping[1] | Relatedness effects controlled? [2] | Age effects controlled?[3] | Form of Help[4] | Notes | Refs |
|---|---|---|---|---|---|---|
| *Sialia mexicana* | Male-biased | Yes | Yes | PPO | Helping data for juveniles still resident within their natal groups. This sex difference in the probability of helping was also apparent for adults within the study area that had the opportunity to help both of their parents. | [1] |
| *Mimus parvulus* | Male-biased | Yes | Yes | PPO | Analyses are of the % of 'potential helpers' that helped. Table 2 confirms male-biased helping is evident within adults of the same age and the text confirms that relatedness to recipients was similar for females and males. While 'potential helpers' included breeders and non-breeders (they were only classed as 'helping' if they fed the offspring of others), non-breeding females were no more likely to 'help' than breeding females, while non-breeding males were significantly more likely to 'help' than breeding males. As such, among non-breeding individuals alone, the sex bias in the incidence of helping should be more male-biased than the significant male-bias already evident among all 'potential helpers'. | [2, 3] |
| *Campylorhynchus nuchalis* | NSB | Yes | Yes | OPR | Sex differences in helper contributions were tested within age and relatedness classes of helpers, and helper contributions did not change with relatedness to resident breeders. | [4, 5] |

| | | | | | | |
|---|---|---|---|---|---|---|
| *Plocepasser mahali* | Female-biased | Yes | Yes | OPR | Strong female bias in helper effort and effects in [6]. [7] verifies female bias still clear when focussing on natal helping only (i.e. in the absence of sex differences in relatedness to recipients) and while controlling for age effects. | [7] |
| *Philetairus socius* | NSB | Yes | Yes | OPR | Data for one year old helpers within their natal groups (as only males help at older ages) | [8] |
| *Acrocephalus sechellensis* | NSB | Yes | Yes | OPR | Table 2 shows no sex difference in the provisioning rates of non-parent subordinates, after allowing for effects of variation in subordinate age and relatedness to recipients. | [9] |
| *Corvus brachyrhynchos* | Female-biased | Yes | Yes | PPO | Sex difference in the probability that yearlings remaining on their natal territory helped to feed young. | [10] |
| *Corvus corone* | Male-biased | Yes | Yes | OPR | Sex difference in the provisioning rates of non-breeding individuals (i.e. helpers) in their natal group, while controlling for (no significant) age effects | [11] |
| *Psilorhinus morio* | Female-biased | Yes | Yes | OPR | Helping data for non-breeding birds analysed controlling for variation in relatedness to the brood, dispersal stage (immigrant vs natal) and age effects. | [12] |
| *Aphelocoma coerulescens* | Male-biased | Yes | Yes | OPR | Significant sex differences apparent both for 'first-year' helpers and 'older' helpers (though sample sizes were small). | [13] |
| *Pomatostomus temporalis* | Female-biased | Yes | Yes | OPR | Helping data for young 'brown eyed' birds. Helpers did not breed at this age in this population [14]. Banded offspring in this population were never observed to disperse before their second year of life, and so these young birds are likely to have been within their natal group. Unpublished genetic analyses mentioned in [14] did suggest, however, that 1 unbanded young bird (i.e. whose dispersal history was not observed) was not within its natal group, leaving it possible that birds of this age do contain rare immigrants. However, as both direct observations and population genetic analyses suggest that this species shows no detectable sex-bias in dispersal ([15]; Table S2), it is unlikely that this sex bias in helping is confounded by a sex difference in the mean relatedness of these young helpers to recipients arising from the inclusion of rare young immigrants, as a similar incidence of these would be expected in both sexes. | [14] |
| *Pomatostomus ruficeps* | NSB | Yes | Yes | OPR | Analysis of the contribution of non-breeding helpers, controlling for variation in relatedness to recipients and helper age (yearlings vs older adults). | [16] |
| *Malurus coronatus* | NSB | Yes | Yes | OPR | Follow-up analysis by author A. Peters, using data from this paper and controlling for variation in helper age and relatedness to the brood (same results if restricting to natal helper with r = 0.5 to recipients) | [17] |

| | | | | | | |
|---|---|---|---|---|---|---|
| *Malurus elegans* | NSB | Yes | Yes | OPR | Author L. Brouwer confirmed data set contained only natal helpers. Methods section demonstrates no effect of age on provisioning rates and so the analyses proceed without fitting age as a covariate predictor. | [18] |
| *Manorina melanophrys* | NSB | Yes | No[5] | OPR | Helper provisioning effort assessed while controlling for variation in relatedness to recipients. Age effects *not* controlled and do appear to confound helper sex (females younger on average). We therefore confirmed that removing this species (and the 3 other species in which potential helper age effects may not have been controlled) from the data set did not qualitatively affect the outcome of our analyses[5] | [19] |
| *Merops bullockoides* | NSB | Yes | No[5] | PPO | Data for potential helpers within their natal clan; "There was no sexual bias in the probability of helping for any category of natal potential helper" p510. The analyses did not apparently control for potential effects of variation in helper age, so we confirmed that removing this species (and the other 3 species with this issue) from the data set did not qualitatively affect the outcome of our analyses[5] | [20] |
| *Dacelo novaeguineae* | Male-biased | Yes | Yes | I | Significant male-bias in (non-breeding) helper contributions to incubation evident in Figure 1 and supported by reported model outcomes while controlling for age and relatedness. The same looks to be true for provisioning rates but the figures do not isolate the sex differences for helpers as clearly. | [21] |
| *Melanerpes formicivorus* | NSB | Yes | No[5] | OPR, TBO | Data for non-breeding helpers (and helpers in this species are offspring that have delayed dispersal from their natal group [22]). No significant sex bias for non-breeding helpers in the analyses that used all available data[6]. Analyses do *not* evidently control for age effects, and helper provisioning rates are known to increase with age [23]. We therefore confirmed that removing this species (and the 3 other species in which potential helper age effects may not have been controlled) from the data set did not qualitatively affect the outcome of our analyses[5] | [22] |
| *Suricata suricatta* | Female-biased | Yes | Yes | OPR, BS | Analyses of provisioning by non-breeding subordinates helping within their natal group and controlling for effects of age | [24] |
| *Mungos mungo* | Male-biased | Yes | Yes | OPR, BS | Data for sub-adults (< 1 year old), which are in their natal groups rarely breed | [25] |
| *Hyaena brunnea* | Female-biased | Yes | Yes | OPR | Significant female-biased contributions to helping among non-breeders, and among sub-adults in their natal group. Sample sizes were very small and analysis may not have been conducted on individual rates, so we confirmed that removing this species | [26] |

| | | | | | | |
|---|---|---|---|---|---|---|
| | | | | | did not qualitatively impact the outcomes of our analyses. | |
| *Meriones unguiculatus* | Female-biased | Yes | Yes | OA | Experimental captive study; all helpers were non-breeding juveniles from the previous litter (i.e. within the natal group). | [27] |
| *Microtus ochrogaster* | NSB | Yes | Yes | OA | For juveniles in intact families (i.e. helping in the natal group) | [28, 29] |
| *Rhabdomys pumilio* | NSB | Yes | Yes | OA | Experimental captive study, with age-matched pairs of male and female non-breeding helpers of two ages helping to rear their parents' young (i.e. within the natal group). | [30] |
| *Fukomys damarensis* | Female-biased | Yes | Yes | OC | Captive study. Non-breeding helpers within their natal groups and the analysis controls for age effects. | [31] |
| *Saguinus oedipus* | Male-biased | Yes | Yes | OC | Captive study. Offspring carrying behaviour by non-breeding brothers and sisters of the focal offspring (i.e. natal helping). Both adults and subadults show significant sex differences in percentage time spent carrying (i.e. analyses age controlled). | [32] |
| *Callithrix jacchus* | Male-biased | Yes | No[5] | OC | This finding and the original source studies are cited in [33, 34], but the source studies (research theses [35, 36]) could not be accessed. We therefore confirmed that removing this species from the analysis had no qualitative impact on the outcomes. There was no suggestion that the original analyses had controlled for age effects, so we also confirmed that removing this species (along with the other 3 species in which helper age effects may not have been controlled), did not qualitatively affect the outcomes of our analyses[5] | [35, 36] |
| *Lamprotornis superbus* [7] | Male-biased | Uncertain [7] | N/A | OPR, PPO | | [37] |
| *Climacteris rufus* [7] | NSB | Uncertain [7] | N/A | OPR | | [38] |
| *Phoeniculus purpureus* [7] | NSB | Uncertain [7] | N/A | PIF | | [39] |
| *Struthidea Cinerea* [7] | Mixed[8] | Uncertain [7] | N/A | OPR | | [40] |
| *Turdoides bicolor* [7] | NSB | Uncertain [7] | N/A | OPR | | [41] |
| *Turdoides caudata* [7] | Male-biased | Uncertain [7] | N/A | PPO | | [42] |
| *Turdoides squamiceps* [7] | NSB | Uncertain [7] | N/A | OPR | Inferred from reported data distributions | [43] |

| | | | | | | |
|---|---|---|---|---|---|---|
| *Canis lupus* [7] | Female-biased | Uncertain [7] | N/A | OA | Further analysis conducted of data presented in the paper | [44] |
| *Helogale parvula* [7] | Female-biased | Uncertain [7] | N/A | BS | Further analysis conducted of data presented in the paper, using observed and expected frequencies of male and female helper contributions | [45] |

[1] 'Female-biased' and 'Male-biased' refer to statistically significant sex differences in helper contributions, while 'NSB' refers to no significant sex bias

[2] 'Yes' where the sex bias in helping was statistically tested for either (i) focusing *solely* on helpers still residing within their natal groups (where helpers are unlikely to differ in their mean relatedness to potential recipients) or (ii) while controlling statistically for the effects of variation in relatedness to recipients on helper effort, if data from helpers in more diverse relatedness contexts were also included in the analysis (e.g. [9, 16, 19]; see species-specific notes for details). Otherwise 'Uncertain [7]' (such species were not included in our analyses [7]).

[3] 'Yes' where the sex bias in helping was tested for among helpers of the same age class or while controlling for effects of variation in helper age (see species-specific notes for details). Otherwise 'No[5]'.

[4] OPR (Offspring provisioning rate), PPO (Probability of provisioning offspring), TBO (% time brooding offspring), I (Incubation), BS (Babysitting), PIF (Provisioning incubating female), OA (Offspring [nest or den] attendance, including licking and grooming), OC (Offspring carrying).

[5] The source study's analysis of the sex bias in helping in these species did *not* evidently control for effects of helper age on helping effort (leaving it possible that the sex difference in dispersal in this species is confounded by a sex differences in mean helper age among the helpers monitored arising from the sex difference in dispersal). We therefore confirmed that the outcomes of our model comparisons were qualitatively unchanged when removing these four species from the analysis. The Dispersal hypothesis model still significantly outperformed the Null (phylogeny only) and Heterogamety models when modelling both female-biased and male-biased helping ($\Delta$AICc > 2 in all cases). And the Dispersal hypothesis model still significantly outperformed both the Parental skills and Paternity uncertainty hypotheses when modelling female-biased helping ($\Delta$AICc > 2 in all cases) but not when modelling male-biased helping (where, once again, none of these three hypothesis models consistently significantly outperformed the null model; $\Delta$AICc < 2 in all cases). As such, our findings cannot be readily attributed instead to the sexes differing in their mean age-at-monitoring within the original studies.

[6] NSB for rates of brooding (Figure 3a in [22]) and provisioning (Figure 3c in [22]) by non-breeding helpers; these figure panels are cited in the text as reflecting the mixed model analyses of all data and show no significant sex differences in helper contributions despite large sample sizes. A Wilcoxon paired-comparison using the subset of nests at which both at least one male and one female non-breeding helper fed, suggested that females helpers fed more than males in this context. But this analysis used a subset of the data and did not account for the repeated measures structure across breeding attempts, while the mixed model analysis of the full data set did (using random factors) and indicated no sex bias. For brooding, a similar paired analysis confirmed the lack of sex-bias in Figure 3a [22].

[7] These species (highlighted in grey) were not included in our analyses as the original source studies tested for a sex difference in helper contributions *without* restricting attention solely to helpers within their natal groups and *without* controlling for variation in helper relatedness to recipients, leaving their outcomes ill-suited to testing the Dispersal hypothesis (as sex differences in dispersal in these species could be confounded by sex differences in mean helper relatedness to recipients within the source analyses; see Methods in the main paper). We include these species here to clarify why they weren't included in our analyses and to show that their sex differences in cooperation (this table) and dispersal (Tables S2) are also consistent with the predictions of the Dispersal hypothesis. As for the 27 species within the analysed set, all of the species that showed significant sex differences in both dispersal and natal cooperation (*Lamprotornis superbus, Turdoides caudata*, *Canis lupus* and *Helogale parvula*) showed the sex difference in natal cooperation that would be predicted by the Dispersal hypothesis on the basis of the sex difference in dispersal. None of the nine species here showed the opposite sex difference in natal cooperation to that which would be predicted by the Dispersal hypothesis on the basis of their sex difference in dispersal.

[8] While this species shows male-biased helper contributions to offspring provisioning, it shows *female*-biased helper contributions to incubation. This doesn't affect our analyses as this species was not included in our analyses (see [7]).

**Table S2 – Data for sex biases in Dispersal and Parenting, and the incidence of Extra-pair and Extra-group paternity**

The first 27 species in the table were used in the analyses presented in the main paper. They constitute the cooperatively breeding bird and mammal species in which both sexes help as non-breeders within their natal group, for which source studies had statistically tested for sex differences in both dispersal (Table S2) and helper contributions within the natal group (or, where helpers of other classes were also included, while controlling for effects of variation in relatedness to recipients). The last 9 species in the table (highlighted in grey) were *not* used in the analyses in the main paper, as they meet the same criteria *except* the source studies' statistical analyses of the sex difference in helper contributions were *not* restricted to helpers within their natal group and did *not* control for variation in helper relatedness to recipients, leaving their outcomes ill-suited to testing the Dispersal hypothesis (see Table S1 legend and footnote 7). 'Female-biased' and 'Male-biased' refer to statistically significant sex differences. 'NSB' refers to no significant sex difference. See table footnotes and Methods for further details.

| Species | Sex-bias in Dispersal | Dispersal Trait [1] | Dispersal References | Sex-bias in Parenting | Parenting References | Extra-pair paternity [2] | Extra-group paternity [3] | Paternity References |
|---|---|---|---|---|---|---|---|---|
| *Sialia mexicana* | Female-biased | T, D | [46] | NSB | [47] | 18.8% | 15.8% | [48] |
| *Mimus parvulus* | Female-biased | I/T | [3] | - | - | - | - | - |
| *Campylorhynchus nuchalis* | Female-biased | I | [4] | NSB [4] | [49] | 10.1% | 1.4% | [50] |
| *Plocepasser mahali* | Male-biased | GS, D | [51] | Female-biased | [52] | 15.0% [5] | 15.0% [5] | [53] |
| *Philetairus socius* | Female-biased | I | [8] | Male-biased | [54] | 0.0% | 0.0% | [55] |
| *Acrocephalus sechellensis* | NSB [6] | I/T | [56] | Female-biased [7] | [57] | 40.0% | 38.2% | [58] |
| *Corvus brachyrhynchos* | Male-biased | I/T | [10] | Male-biased | [59] | 17.3% | 10.4% | [60] |
| *Corvus corone* | Female-biased | I/T | [61] | NSB | [11] | 10.7% [5] | 0.0% | [62] |
| *Psilorhinus morio* | Male-biased | I/T | [63] | NSB | [64] | 80.0% | 24.0% | [65] |
| *Aphelocoma coerulescens* | Female-biased | T, D | [66, 67] | Male-biased | [13] | 0.0% | 0.0% | [68] |
| *Pomatostomus temporalis* | NSB | GS | [15] | - | - | 23.9% [8] | 19.7% [8] | [69] |
| *Pomatostomus ruficeps* | Female-biased | I, GS | [70] | Male-biased | [71] | 8.0% | 0.0% | A.F.Russell pers. comm. |

| | | | | | | | | |
|---|---|---|---|---|---|---|---|---|
| *Malurus coronatus* | Female-biased | I/T, D | [72, 73] | NSB | [74] | 4.4% | 2.6% | [75] |
| *Malurus elegans* | Female-biased | I, T | [76] | NSB | [18] | 57.0% | 55.1% | [77] |
| *Manorina melanophrys* | Female-biased | I/T | [78] | Female-biased | [19] | 4.2% | 4.2% | [79] |
| *Merops bullockoides* | Female-biased | I | [20] | - | - | 4.1% [5] | 2.6% [5] | [80] |
| *Dacelo novaeguineae* | Female-biased | I/T | [81] | Male-biased | [21] | 0.0% | 0.0% | [81] |
| *Melanerpes formicivorus* | Female-biased | I, D | [82] | Female-biased | [22] | JNP [9] | JNP [9] | - |
| *Suricata suricatta* | Male-biased | I | [24, 83] | Female-biased | [84] | 8.0% [8] | 5.0% [8] | [85] |
| *Mungos mungo* | NSB | I/T | [86] and M.A.Cant pers. comm. | - | - | JNP [9] | JNP [9] | - |
| *Hyaena brunnea* | Male-biased | GS | [87] | - | - | - | - | - |
| *Meriones unguiculatus* | Male-biased | GS | [88] | Female-biased | [27] [10] | JNP [9] | JNP [9] | - |
| *Microtus ochrogaster* | Male-biased | I, GS | [89-92] [11] | Female-biased | [29] | JNP [9] | JNP [9] | - |
| *Rhabdomys pumilio* | Male-biased | GS | [93] | Female-biased | [30] | JNP [9] | JNP [9] | - |
| *Fukomys damarensis* | Male-biased | GS | [94] | - | - | 18.1% | 10.5% | [95] |
| *Saguinus oedipus* | NSB | I/T | [96] | Male-biased | [32] [10] | - | - | - |
| *Callithrix jacchus* | Female-biased | I | [97] | NSB | [98] [10] | - | - | - |
| *Lamprotornis superbus* [12] | Female-biased | I | [99] | | | | | |
| *Climacteris rufus* [12] | Female-biased | I/T | [38] | | | | | |
| *Phoeniculus purpureus* [12] | NSB | I | [100] | | | | | |

| | | | |
|---|---|---|---|
| *Struthidea cinerea* [12] | NSB | GS | [101] |
| *Turdoides bicolor* [12] | NSB | I, T, GS | [102, 103] |
| *Turdoides caudata* [12] | Female-biased | I/T | [42] |
| *Turdoides squamiceps* [12] | Female-biased | I | [104] |
| *Canis lupus* [12] | Male-biased | I, GS | [105, 106] [13] |
| *Helogale parvula* [12] | Male-biased | I/T | [107] |

[1]The dispersal traits used to determine sex differences in dispersal: I (dispersal incidence), T (dispersal timing), I/T (dispersal incidence or timing; it can be unclear which is driving a given reported pattern), GS (genetic structure), D (dispersal distance).

[2]The proportion of offspring not sired by the dominant breeding male. Where possible this value was calculated for the broods or litters of the dominant/primary female (the scenario envisaged by the *Paternity uncertainty hypothesis*).

[3]The proportion of offspring not sired by a male resident within the group. Where possible this value was calculated for the broods or litters of the dominant/primary female (the scenario envisaged by the *Paternity uncertainty hypothesis*).

[4]No statistical analysis presented, but evident from Figure 10 in the source paper.

[5]Average of the highest and lowest possible population-level values for %EPP or %EGP, given residual uncertainty following assignments.

[6]An initial male bias in dispersal disappeared over time, and has since been attributed to the overproduction of females on higher quality territories (from which offspring of both sexes are more likely to delay dispersal).

[7]No statistical analysis presented, but evident from Figure 6 in the source paper.

[8]Using only the paternity figures provided for the broods or litters of the dominant/primary female (the scenario envisaged by the Paternity uncertainty hypothesis).

[9]Joint-nesting plural breeder (JPN) – these species were excluded from Paternity uncertainty hypothesis model tests given the expectation of maternity uncertainty among females (which would void a key assumption of the Paternity uncertainty hypothesis).

[10]Captive study.

[11]NSB sometimes reported [92, 108]. Fitting this species as NSB for dispersal (rather than male-biased) did not qualitatively affect the outcome of our analyses (the species is also NSB for natal helping; Table S1).

[12]See footnote 7 in Table S1

[13]NSB sometimes reported [109, 110].

**Table S3 - Species in our dataset not present in the source mammal and bird supertrees.** For six species it was possible to utilize the position of a closely related species in the same genus, thereby maintaining the same relative phylogenetic relationships between species. For two other species it was necessary to add new branches with appropriate sister-taxa relationships with the other species in our analysis (see notes in the table).

| Species with data | Species used in tree | Notes |
|---|---|---|
| *Hyaena brunnea* | *Hyaena hyaena* | Same genus |
| *Meriones unguiculatus* | *Meriones dahli* | Same genus |
| *Pomatostomus temporalis* | *Pomatostomus superciliosus* | Same genus |
| *Corvus brachyrhynchos* | *Corvus macrorhynchos* | Same genus |
| *Malurus coronatus* | *Malurus splendens* | Same genus |
| *Mimus parvulus* | *Mimus thenca* | Same genus |
| *Plocepasser mahali* | New branch | Classified with the true weavers (*Ploceidae*; formerly classified with the true sparrows, *Passeridae*). Our sample contains one other weaver species (*Philetairus socius*) and so *P. mahali* was positioned as a sister-taxon to *P. Socius* in our tree (see Figure 1 main paper) |
| *Psilorhinus morio* | New branch | Ericson et al. [111] show *Psilorhinus* and *Aphelocoma* as being more closely related to each other than either is to the other genus of *Corvidae* used in this analysis (*Corvus*). *Psilorhinus morio* was therefore positioned as a sister-taxon to *Aphelocoma coerulescens* in our tree (see Figure 1 main paper) |

**Table S4 – Impacts of branch length back to the avian/mammalian common ancestor** The impact of different assumptions about the scaling of the branch lengths (BL) joining mammals and birds to their common ancestor (see Figure S1 and methods) on the strength of support for the sex bias in dispersal (MB, NSB, FB) predicting the probability of (i) Female-biased helping (vs Other) and (ii) Male-biased helping (vs Other), in our phylogenetic logistic regressions. The branches joining mammals and birds were scaled as either being equal to all other branches (the assumption used for the analyses presented in the main text), or 5, 10 or 100 times longer than the other branches. Changing this assumption from that utilized in the results reported in the main paper (BL=1), only increased the strength of ΔAICc support for the dispersal hypothesis model relative to the null (phylogeny only) model in the model of Male-biased helping, suggesting that the approach applied in the main paper is likely to be conservative. Doing so had no effect on the strength of ΔAICc support in the model of Female-biased helping because the phylogenetic signal was much weaker in this model.

| | Modeling the probability of **Female-biased helping** | Modeling the probability of **Male-biased helping** |
|---|---|---|
| Branch length scaling factor | ΔAICc of Dispersal model vs Null (phylogeny only) model | ΔAICc of Dispersal model vs Null (phylogeny only) model |
| 1 | 13.40 | 3.53 |
| 5 | 13.40 | 4.40 |
| 10 | 13.40 | 4.78 |
| 100 | 13.40 | 5.20 |

| Predictor | Species | BL=1 | Rho=1 | Rho=2 | Rho=3 | Lambda=0 | MPLE | Standard LR | $R^2$ range |
|---|---|---|---|---|---|---|---|---|---|
| **Comparing explanatory power of Sex Bias (SB) in Dispersal vs Heterogamety & the Null model** | | | | | | | | | |
| Null model | N = 27 | **37.32** | 37.32 | 37.32 | 37.32 | 37.32 | 37.32 | 34.95 | - |
| SB in Dispersal (FB, NSB, MB) | N = 27 | **23.93** | 23.79 | 23.69 | 23.60 | 23.93 | 24.89 | 20.87 | 0.54-0.65 |
| Heterogamety (mammal vs bird) | N = 27 | **38.50** | 38.50 | 38.50 | 38.50 | 38.50 | 38.51 | 35.84 | 0.05 |
| **Comparing explanatory power of Sex Bias (SB) in Dispersal vs Sex Bias in Parental Care & the Null model** | | | | | | | | | |
| Null model | N = 21 | **27.74** | 27.74 | 27.74 | 27.74 | 27.74 | 27.75 | 25.24 | - |
| SB in Dispersal (FB, NSB, MB) | N = 21 | **19.70** | 19.70 | 19.70 | 19.70 | 20.56 | 20.57 | 15.64 | 0.55-0.62 |
| SB in Parental Care (FB, NSB, MB) | N = 21 | **32.42** | 32.42 | 32.42 | 32.44 | 32.40 | 32.45 | 28.99 | 0.06 |
| **Comparing explanatory power of Sex Bias (SB) in Dispersal vs Extra-Pair / Extra-Group Paternity & the Null model** | | | | | | | | | |
| Null model | N = 18 | **27.72** | 27.72 | 27.72 | 27.72 | 27.72 | 27.72 | 25.11 | - |
| SB in Dispersal (FB, NSB, MB) | N = 18 | **13.85** | 13.85 | 13.85 | 13.85 | 13.85 | 15.66 | 10.04 | 0.77-0.88 |
| Extra-Pair Paternity | N = 18 | **26.77** | 26.40 | 26.35 | 26.37 | 26.77 | 26.78 | 23.47 | 0.16-0.17 |
| Extra-Group Paternity | N = 18 | **26.39** | 24.81 | 24.19 | 24.10 | 26.57 | 26.45 | 23.33 | 0.18-0.20 |

**Table S5 - Comparison of alternative hypotheses to explain the probability of FEMALE biased helping, using different phylogenetic assumptions.** The rows capture the competing hypotheses (and their key predictor). 'Null model' is the phylogeny-only model. 'SB in Philopatry' is a model that includes the sex bias in philopatry (FB, NSB, MB) as the only predictor (alongside phylogeny). 'Heterogamety' is a model that includes whether the species is a bird or a mammal as the only predictor (alongside phylogeny). 'SB in Parental Care' is a model that includes the sex bias in parental care (FB, NSB, MB) as the only predictor (alongside phylogeny). "Extra-pair [or group] paternity" is a model that includes the incidence of extra-pair [or group] paternity as the only predictor (alongside phylogeny). The first group of three rows presents model comparisons using the full data set of 27 species for which data was available for testing the three hypotheses considered therein. The second group of three rows presents model comparisons using the 21 species with sex bias in parental care information available. The final group of four rows presents model comparisons using the 18 species with extra-pair (or group) paternity information. The different columns between the 'Species' column and '$R^2$ range' columns, present the AICc support for the focal model under different sets of phylogenetic assumptions (lower values reflect stronger support and a ΔAICc >2 between any pair of models utilizing the same data set can be interpreted as reflecting stronger statistical support for the model with the lower AICc value). First, we test different assumptions regarding branch lengths when using the Ives & Garland [112] approach to phylogenetic logistic regression: (i) BL = 1, all branch lengths are assumed to be 1 (the results reported in the main paper; and hence highlighted in bold here); (ii) scaling branch lengths via Rho = 1, 2 and 3 (see methods; Figure S2); and (iii) Lambda = 0: which is equivalent to assuming that there is no phylogenetic signal. 'MPLE' presents the findings using the Maximum Penalized Likelihood Estimation methodology (with BL=1) in place of the Ives & Garland [112] methodology. 'Standard LR' presents the outcomes under standard logistic regression (i.e. not controlling for phylogeny). '$R^2$ range': we use Tjur's method [113] for calculating a psuedo R-squared value for logistic regression.

| Predictor | Species | BL=1 | Rho=1 | Rho=2 | Rho=3 | Lambda=0 | MPLE | Standard LR | $R^2$ range |
|---|---|---|---|---|---|---|---|---|---|
| **Comparing explanatory power of Sex Bias (SB) in Dispersal vs Heterogamety** | | | | | | | | | |
| Null model | N = 27 | **36.95** | 37.05 | 37.02 | 37.02 | 37.32 | 36.98 | 34.95 | - |
| SB in Dispersal (FB, NSB, MB) | N = 27 | **33.42** | 33.17 | 33.09 | 32.97 | 34.52 | 34.38 | 31.50 | 0.21-0.26 |
| Heterogamety (mammal vs bird) | N = 27 | **39.50** | 39.59 | 39.57 | 39.57 | 39.79 | 39.53 | 37.13 | 0.00 |
| **Comparing explanatory power of Sex Bias (SB) in Dispersal vs Sex Bias in Parental Care** | | | | | | | | | |
| Null model | N = 21 | **29.46** | 29.80 | 29.80 | 29.81 | 29.80 | 29.48 | 27.30 | - |
| SB in Dispersal (FB, NSB, MB) | N = 21 | **29.00** | 29.30 | 28.62 | 28.19 | 29.62 | 29.85 | 26.21 | 0.19-0.24 |
| SB in Parental Care (FB, NSB, MB) | N = 21 | **28.42** | 28.42 | 27.92 | 27.55 | 28.42 | 29.29 | 25.01 | 0.22-0.32 |
| **Comparing explanatory power of Sex Bias (SB) in Dispersal vs Extra-Pair (or Extra-Group) Paternity** | | | | | | | | | |
| Null model | N = 18 | **23.89** | 23.89 | 23.89 | 23.89 | 23.89 | 23.89 | 21.30 | - |
| SB in Dispersal (FB, NSB, MB) | N = 18 | **27.47** | 27.12 | 28.18 | 28.16 | 27.11 | 27.09 | 22.01 | 0.13-0.18 |
| Extra-Pair Paternity | N = 18 | **24.15** | 24.15 | 24.15 | 24.15 | 24.23 | 24.15 | 21.07 | 0.14-0.15 |
| Extra-Group Paternity | N = 18 | **23.62** | 78.55 | 23.62 | 23.62 | 23.62 | 23.60 | 20.46 | 0.09-0.18 |

**Table S6 - Comparison of alternative hypotheses to explain the probability of MALE biased helping, using different phylogenetic assumptions.** The rows capture the competing hypotheses (and their key predictor). 'Null model' is the phylogeny-only model. 'SB in Philopatry' is a model that includes the sex bias in philopatry (FB, NSB, MB) as the only predictor (alongside phylogeny). 'Heterogamety' is a model that includes whether the species is a bird or a mammal as the only predictor (alongside phylogeny). 'SB in Parental Care' is a model that includes the sex bias in parental care (FB, NSB, MB) as the only predictor (alongside phylogeny). "Extra-pair [or group] paternity" is a model that includes the incidence of extra-pair [or group] paternity as the only predictor (alongside phylogeny). The first group of three rows presents model comparisons using the full data set of 27 species for which data was available for testing the three hypotheses considered therein. The second group of three rows presents model comparisons using the 21 species with sex bias in parental care information available. The final group of four rows presents model comparisons using the 18 species with extra-pair (or group) paternity information. The different columns between the 'Species' column and '$R^2$ range' columns, present the AICc support for the focal model under different sets of phylogenetic assumptions (lower values reflect stronger support and a ΔAICc >2 between any pair of models utilizing the same data set can be interpreted as reflecting stronger statistical support for the model with the lower AICc value). First, we test different assumptions regarding branch lengths when using the Ives & Garland [112] approach to phylogenetic logistic regression: (i) BL = 1, all branch lengths are assumed to be 1 (the results reported in the main paper; and hence highlighted in bold here); (ii) scaling branch lengths via Rho = 1, 2 and 3 (see methods; Figure S2)*;* and (iii) Lambda = 0: which is equivalent to assuming that there is no phylogenetic signal. 'MPLE' presents the findings using the Maximum Penalized Likelihood Estimation methodology (with BL=1) in place of the Ives & Garland [112] methodology. 'Standard LR' presents the outcomes under standard logistic regression (i.e. not controlling for phylogeny). '$R^2$ range': we use Tjur's method [113] for calculating a psuedo R-squared value for logistic regression.

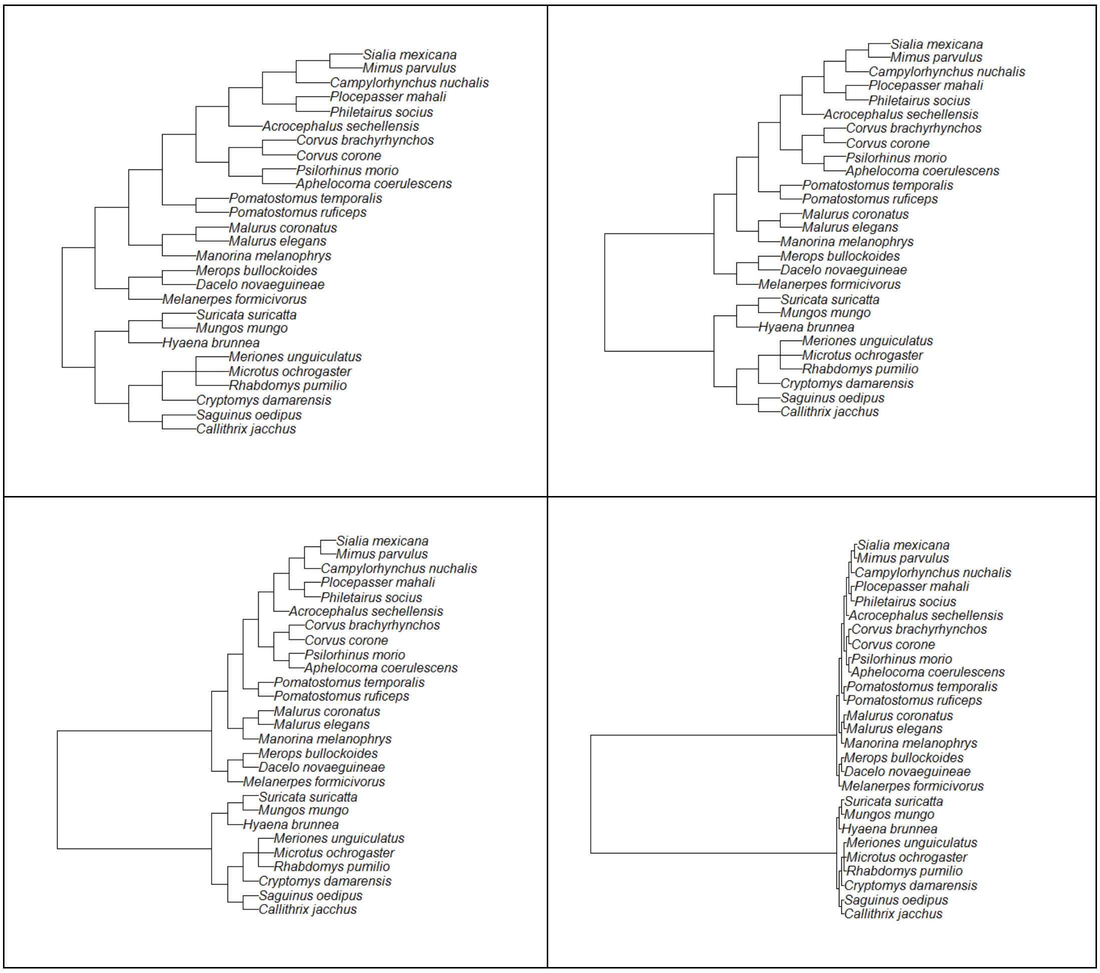

**Figure S1 - Visualization of changing the assumption about the length of the branches joining mammals and birds.** The equal branch length assumption (Top left; as utilized in the analyses presented in the main text) is compared to the branch joining mammals and birds being 5 times (Top right), 10 times (Bottom left), or 100 times (Bottom right) as long as the other branches.

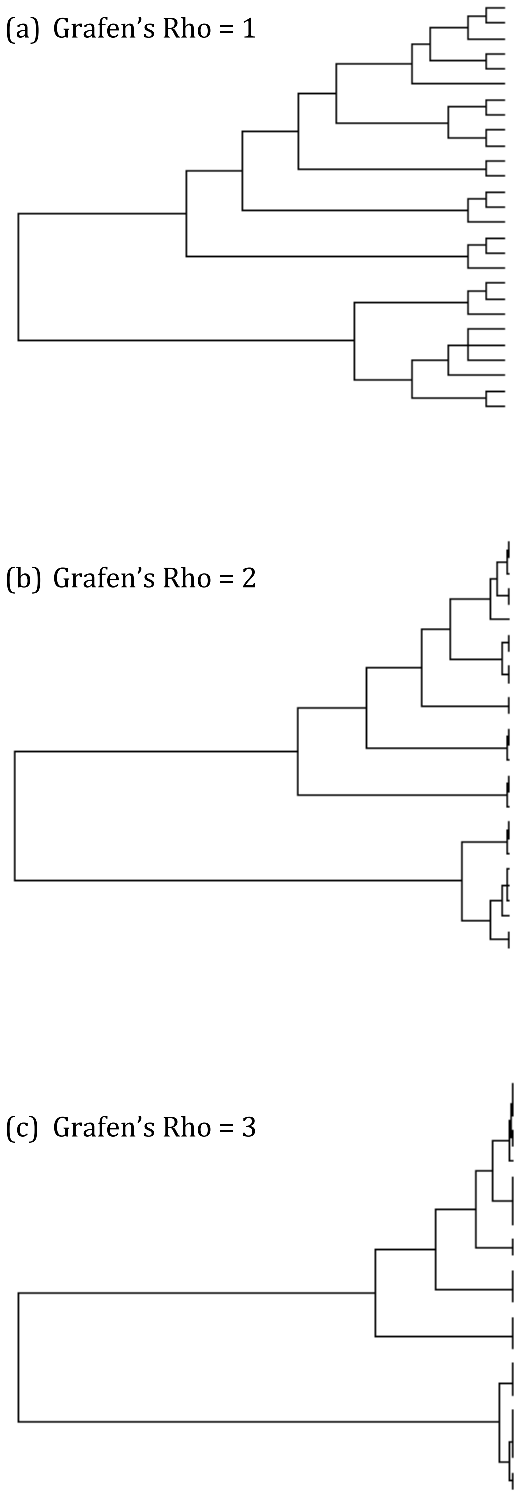

**Figure S2 - Visualization of the effect of Grafen's scaling parameter (Rho) [114] on the relative lengths of branches in the phylogenetic tree**. The algorithm stretches the branches so that the tree is ultrametric (i.e. the root to tip length is the same for all tips) based on the number of nodes descending from each node in the tree. When Rho=1 the scaling is linear throughout the tree from root-to-tip, whereas higher values of Rho result in relatively longer branch lengths near the root of the tree and relatively shorter branch lengths near the tips.

(a)

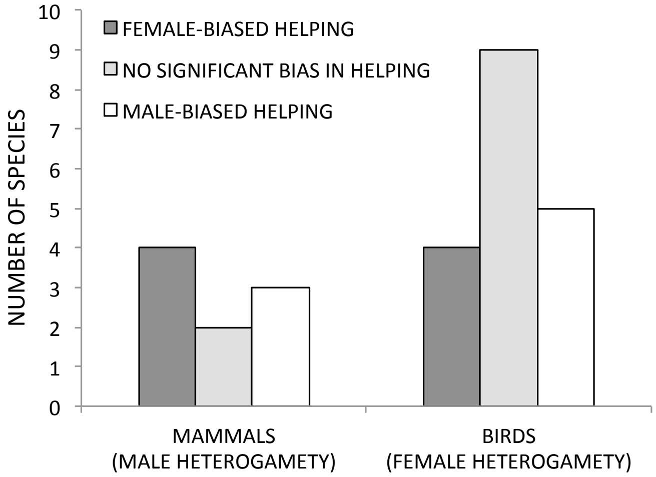

(b)

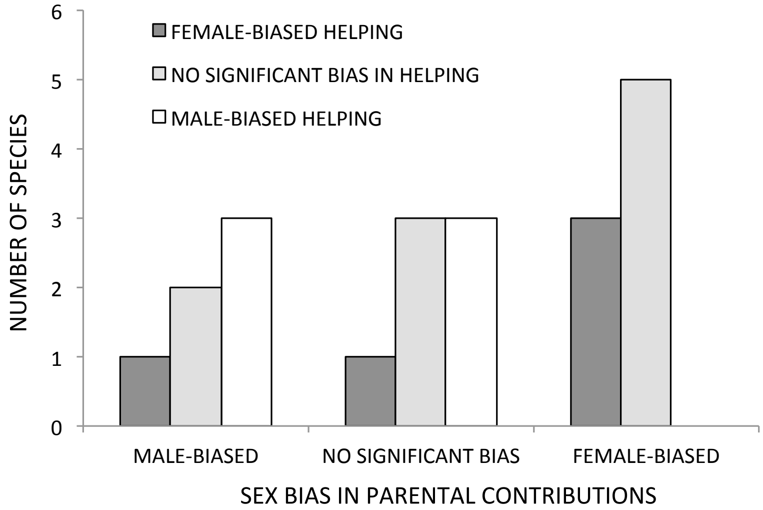

(c)

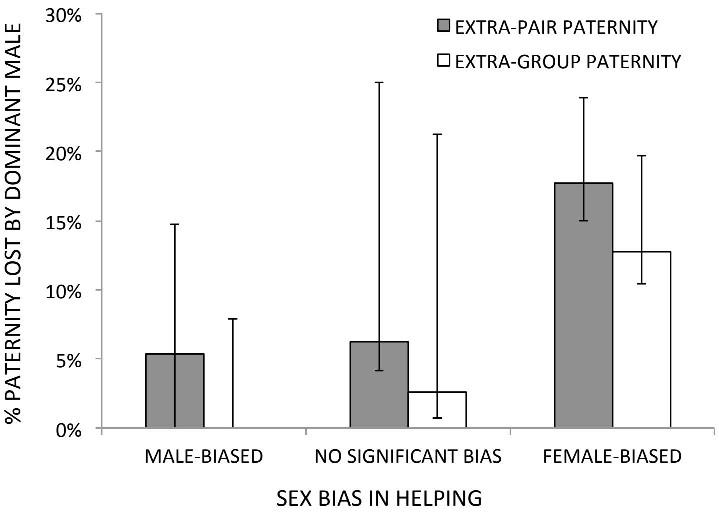

**Figure S3 – The raw data patterns relevant to the three alternative hypotheses, for reference**. Note that these are the patterns within the data set of species collated here specifically for testing the Dispersal hypothesis. Attempts to formally test each of these alternative hypotheses in isolation (not the goal of this study) could therefore utilize data from additional species that might be ill-suited to testing the Dispersal hypothesis. **(a) The Heterogamety hypothesis** predicts that the heterogametic sex (males in mammals and females in birds) should contribute less to helping within its natal group. Whether a species was a mammal or a bird did not predict the nature of its sex bias in helping in this data set (n = 27 species in the full data set; see Tables S5&6 for phylogenetic model comparisons; Fisher's 3x2 exact test without phylogenetic control p = 0.45). **(b) The Parental skills hypothesis** predicts that the sex bias in parental contributions to a given form of parental care should predict the sex bias in helper contributions to that same form of care. This was not the case in this data set (n = 21 species [the subset of the overall 27 for which sufficient parental care data were available]; see Tables S5&6 for phylogenetic model comparisons; Fisher's 3x3 exact test without phylogenetic control p = 0.23). **(c) The Paternity uncertainty hypothesis** predicts that species with a higher incidence of extra-pair (or extra-group) paternity should be more likely to show male-biased helping within the natal group (bars present medians ± IQRs). In this data set, a species' sex bias in helping was not significantly associated with the extent to which dominant males lost paternity to either (i) any other males (i.e. the incidence of extra-pair paternity; n = 18 species with paternity data available; see Tables S5&6 for phylogenetic model comparisons; Kruskal-Wallis test without phylogenetic control: H = 3.51, p = 0.17) or (ii) specifically extra-group males (i.e. the incidence of extra-group paternity; n = 18 species with paternity data available; Tables S5&6; H = 4.52, p = 0.10). These modest p values reflect a weak trend running *counter* to the Paternity uncertainty hypothesis prediction: if anything, dominant males tended to lose more paternity in species with female-biased helping.

### SUPPLEMENTAL DISCUSSION

The lack of support for the Heterogamety, Parental skills and Paternity uncertainty hypotheses in our analyses suggests that they do not provide credible alternative explanations for our findings in support of the Dispersal hypothesis (the focal hypothesis under test here) and are unlikely to be the *primary* driver of sex differences in cooperation in the context studied here. While the mechanisms envisaged in these hypotheses could nevertheless still have contributed to selection for sex differences in cooperation in this and other contexts (see below), there are plausible biological explanations for them not predicting the sex differences in natal cooperation studied here. The lack of any clear effect of heterogamety, which was predicted to impact sex biases in cooperation through sex differences in relatedness for genes sited on the sex chromosomes [115], is consistent with arguments that such effects may be constrained in reality by intra-genomic conflict with genes on the autosomes, and the impact of dosage compensation mechanisms on the link between genotypes and phenotype [116]. The lack of support for the Parental skills hypothesis [117] is consistent with the lack of compelling support to date for its key assumption that helping enhances parental skills [23, 118-120]. And the lack of support for the Paternity uncertainty hypothesis [121] could reflect complexities not captured by the scenario envisaged in the original model. We sought to minimize such complexity when testing the hypothesis by (i) excluding joint-nesting species (in which maternity may also be uncertain), and (ii) testing for a potential effect of extra-group as well as extra-pair paternity, as the former could be the more appropriate predictor if males lose paternity to within-group relatives. Nevertheless, numerous factors could still complicate the impact of lost paternity on patterns of helping. For example, where paternity is lost in part to helper males in other groups [e.g. 122], investment in forays to secure this paternity could leave male helpers in species with extra-group paternity contributing less to cooperative care than female helpers [due to trade-offs between mating forays and natal cooperation; 123], counter to the prediction of the Paternity uncertainty hypothesis.

While our analyses revealed no support for the Heterogamety, Parental skills and Paternity uncertainty hypotheses, this outcome should not be interpreted as evidence that these hypotheses have played no role in selection for sex differences in cooperation. First, our analyses had modest sample sizes, given the need to restrict attention to data collected in the specific contexts required to test the Dispersal hypothesis. While these sample sizes were sufficient to yield statistical support for the Dispersal hypothesis, they may have been insufficient to reveal the potentially more subtle effects of these other hypotheses. Second, our analyses did not test for evidence that these other hypotheses have played a secondary role in shaping sex differences in natal cooperation, acting alongside the Dispersal hypothesis, because our modest sample sizes leave our data set ill-suited to simultaneously estimating the effects of multiple predictors alongside the phylogeny. Third, our analyses focused on the species to which the rationale of the Dispersal hypothesis could apply: species in which both sexes delay dispersal and help within their natal group. While these other hypotheses may not be the primary driver of sex differences in cooperation in this context, they could conceivably play a role in other contexts. For example, in some cooperative breeders only one sex delays dispersal and so only one sex is available to help within the natal group [119, 124-126]. Such species were excluded from our analyses (as the sex difference in dispersal necessarily determines the sex difference in natal helping in such species), but explaining this separate, yet similarly enigmatic, aspect of sex differences in cooperation remains an outstanding challenge [119]. The answer to this riddle likely lies in patterns of selection for delayed dispersal and/or parental toleration of it, but any mechanisms that generate sex differences in the direct payoff from natal cooperation (including those envisaged in these other hypotheses) could conceivably play a role by impacting the fitness payoff available were offspring to delay dispersal and help [120, 127].

Our analyses provide support for the Dispersal hypothesis, which recognizes that sex differences in dispersal have the potential to drive sex differences in natal cooperation via two general mechanisms (that could act in isolation or in concert in any given species): the more dispersive sex may (i) stand to gain a lower direct fitness *benefit* from natal helping [24, 128] and/or (ii) experience a greater direct fitness *cost* of natal helping [12, 123]. We suggest that variation among species in the relevance of these two potential mechanisms could help to explain deviations within our data set from the patterns predicted by the Dispersal hypothesis. For example, ten species showed significant sex biases in dispersal without a significant sex bias in natal helping (Figure 2a). These deviations from the predicted pattern could reflect limited statistical power in certain helping studies, but it also seems likely that in some species the less dispersive sex may not actually stand to gain a greater net direct fitness payoff from helping (because the mechanisms by which such a link could occur need not apply in all species), in which case no dispersal-related sex bias in natal helping effort would be predicted. For example, all eight species with significantly female-biased dispersal but no significant sex bias in natal helper contributions are birds, in which flight could conceivably ameliorate dispersal-driven sex differences in the direct payoffs from helping in some species, by (i) reducing the downstream direct benefits to be accrued from helping to augment group size (e.g. if group size has a lesser effect on survival where flight facilitates predator evasion), and/or (ii) ameliorating the trade-off between helping and seeking dispersal opportunities (as flight will likely facilitate prospecting). More generally, the extent to which sex biases in dispersal promote sex biases in natal cooperation may depend upon the nature of the benefits and costs of cooperation in any given species, as these could well be more sensitive to dispersal patterns in some species than others. Indeed, while the Dispersal hypothesis rationale emphasizes the potential role of trade-offs between investments in cooperation and dispersal, trade-offs between cooperation and other fitness-enhancing traits could explain the handful of species in our analyses that show sex-biased helping without an evident sex-bias in dispersal. For example, the banded mongoose, *Mungos mungo*, shows significantly male-biased contributions to natal helping in the absence of a clear sex bias in dispersal (Figure 1); a finding that has been attributed to trade-offs between helping and future reproduction being stronger in females than males [25].
